# Importance of test-retest reliability for promoting fMRI based screening and interventions in major depressive disorder

**DOI:** 10.1101/2020.12.11.421750

**Authors:** Laurie Compère, Greg J. Siegle, Kymberly Young

## Abstract

Proponents of personalized medicine have promoted neuroimaging evaluation and treatment of major depressive disorder in three areas of clinical application: clinical prediction, outcome evaluation, and neurofeedback. Whereas psychometric considerations such as test-retest reliability are basic precursors to clinical adoption for most clinical instruments, they are often not considered for neuroimaging assessments. As an example, we consider functional magnetic resonance imaging (fMRI) of depression, a common and particularly well validated mechanistic technology for understanding disorder and guiding treatment. In this article, we review work on test-retest reliability for depression fMRI studies. We find that basic psychometrics have not been regularly attended to in this domain. For instance, no fMRI neurofeedback study has included measures of test-retest reliability despite the implicit assumption that brain signals are stable enough to train. We consider several factors that could be useful to aid clinical translation including 1) attending to how the BOLD response is parameterized, 2) identifying and promoting regions or voxels with stronger psychometric properties 3) accounting for within-individual changes (e.g., in symptomatology) across time and 4) focusing on tasks and clinical populations that are relevant for the intended clinical application. We apply these principles to published prognostic and neurofeedback data sets. The broad implication of this work is that attention to psychometrics is important for clinical adoption of mechanistic assessment, is feasible, and may improve the underlying science.

## 1. GENERAL INTRODUCTION

Proponents of personalized medicine have promoted mechanistic evaluation and mechanistically targeted treatments for major depressive disorder (Hansen and Siegle, 2015). As an example, we consider functional magnetic resonance imaging (fMRI), a common and particularly well validated mechanistic technology that represents a promising proof-of-concept in this area. Longitudinal assessment of changes in regional brain activity using functional magnetic resonance imaging (fMRI) has increasingly been used in research on the treatment of psychiatric conditions including major depressive disorder (MDD) (Fournier et al., 2014). As good psychometric properties are essential for any measure to be considered for clinical adoption (Pickford and Guilford, 2007), best-practice guidelines for increasing generalizability and reproducibility of fMRI results are emerging (Nichols et al., 2017; Poldrack et al., 2017). We focus here on test-retest reliability in task-based fMRI and neurofeedback (fMRI-nf) designs, using MDD as a running case example. Ideally, our observations can be applied to other technologies and across neuropsychiatric disorders.

To understand the current state of the field, we conducted literature reviews quantifying how often test-retest reliability was reported in fMRI biomarker and real-time fMRI neurofeedback (rtfMRI-nf) studies in MDD. As we will demonstrate below, this was infrequent and the general literature has shown that wen assessed, reliability was generally low. We thus suggest a few analytic techniques for improving test-rest reliability in fMRI and its clinical applicability. We focus on data analysis to make our suggestions maximally applicable to already collected data. Finally, we test these suggested principles on published MDD neuroimaging treatment outcome and neurofeedback datasets as proofs of concept.

The idea that fMRI could have therapeutic utility is based on assumptions that hemodynamic activity is reliable over time in the absence of intervention, and that observed changes between one scan session and the next have significant and interpretable values (Barch and Mathalon, 2011). The reliability of fMRI also affects its criterion validity, as poor reliability limits the strength of association between the instrument and other relevant measures (Vul et al., 2009).

### 1.1. On computing reliability of fMRI

Demonstrating ability to achieve similar results over time, or the reliability of measures is considered critical to creating a clinically useful measure (Pickford and Guilford, 2007). Reliability is a quantitative measure of stability of an individual’s data (Bennett and Miller, 2013). It refers to the ability of a measure to distinguish participants from each other and to replicate the order of individuals’ ranks during repeated assessments, assuming they do not experience true signal change between assessments (Barch and Mathalon, 2011).

Though stable regional hemodynamic activations at the group level can be observed over time, there are significant changes in how each subject contributes individually to the observed group activation (Caceres et al., 2009; Zandbelt et al., 2008). Various approaches have been used to measure test-retest reliability for fMRI. For example, a Pearson correlation between visits across time measure the degree to which visits on two occasions are linearly related, where data from each visit are independently scaled (e.g., Harrington et al., 2006). A more common approach, and the measure we focus on in this manuscript, involves computing intra-class correlation coefficients (ICC) that also reflect rank ordering of values across days (Bennett and Miller, 2010) as a ratio of variance between values observed across subjects and sites divided by the total visit variance (Bartko, 1966). Values range from 0 (no reliability) to 1 (perfect reliability). There are three different types of ICCs described by the princeps article written by Shrout and Fleiss (1979). The ICC(1,1) index is similar to the Pearson correlation but normalizes by the pooled mean and variance across visits. ICC(2,1) is an agreement index that allows generalization of results across scanners while ICC(3,1) works under the assumption that the variance is the same across scanners. Therefore, the ICC(3,1), mostly used across studies, is a scanner consistency index where the effect of scanner is considered a fixed effect (Shrout and Fleiss, 1979). In order to match the literature in the field and because we considered the scanner as a covariate of interest when investigating the impact of taking into account clinical and design covariates when computing reliability indexes, we mainly used ICC(3,1) in our analyses.

Interpretation of ICC values is subjective with no uniformly accepted standards; ICC values of less than 0.4 are often considered poor, 0.4-0.59 fair, 0.60-0.74 good, and above 0.75 excellent (Cicchetti, 1994; Plichta et al., 2012; Shrout and Fleiss, 1979), though more stringent cutoffs have also been recommended (e.g., Portney and Watkins, 2009). Negative ICC value are usually interpreted as no reliability (Bartko, 1976), since these values are outside the theoretical limits of ICC (although negative values may appear when within-subject variance is greater than between-subject variance) (Lahey et al., 1983).

Though the ICC has been recommended for use in fMRI (Caceres et al., 2009), some fMRI analysis packages (SPM, FSL) do not inherently support computation of this metric, potentially hinting at its perceived value in the field, though other packages (e.g., AFNI, NIfTI) do provide for its computation, and add-on packages (e.g., reliability toolbox for SPM or other packages on R) do allow such computations (see Computation of voxelwise ICCs using different tools in Box 3 in supplementary materials for more details). Indeed, reliability estimates have been rarely reported in fMRI studies and usually reveal poor reliability when estimated (Elliott et al., 2020). Non-clinical studies have generally found low to moderate test-retest reliability values for regional fMRI activity, with ICCs ranging from 0.33-0.66 (reviewed in Bennett and Miller, 2010).

### 1.2. Biomarker/Prediction Studies Review

Many studies suggest fMRI measurements can be used to predict treatment outcome in MDD (for reviews, see Arnone, 2019; Fonseka et al., 2018; Phillips and Swartz, 2014; Wessa and Lois, 2015). The underlying assumption is that biomarkers in the brain are involved in the causal process of MDD. Therefore, it is expected that the activity measured in these biomarkers is related to, and evolves over time with, symptom changes in general and that for interventions targeting the biomarker the more abnormal activity observed, the more effective the intervention will be. However, clinical applications of these findings are limited by the possibility that these biomarkers may have low test-retest reliability (Nord et al., 2017). If a biomarker is not reliable, it is impractical to interpret its activation at the individual level (Fu et al., 2013; Guo et al., 2012). Thus, despite strong predictive utility, researchers acknowledge that their results might be limited by poor test-retest reliability (e.g., Fu et al., 2015). Of particular interest, the amygdala, a commonly reported biomarker for MDD, shows poor to good reliability when emotional stimuli are displayed, with great heterogeneity between studies in healthy participants (Lois et al., 2018). Thus, we surveyed the predictive fMRI literature in MDD to examine whether this first step was being taken.

#### 1.2.1. Method

A PubMed search with the key words “fMRI AND biomarker OR prediction OR predict AND depression OR MDD OR major depressive disorder NOT Rest NOT Resting” produced 140,640 results. We combined this list with other articles discovered in our submitted fMRI meta-analysis of depression treatment outcome prediction studies (Strege et al., 2020) to complete the list of articles (Table 1).” After removing articles not including functional neuroimaging (i.e., studies focusing on volumetric measures or using PET) or human participants, we were left with 55 studies (Table 1).

**Table 1:**
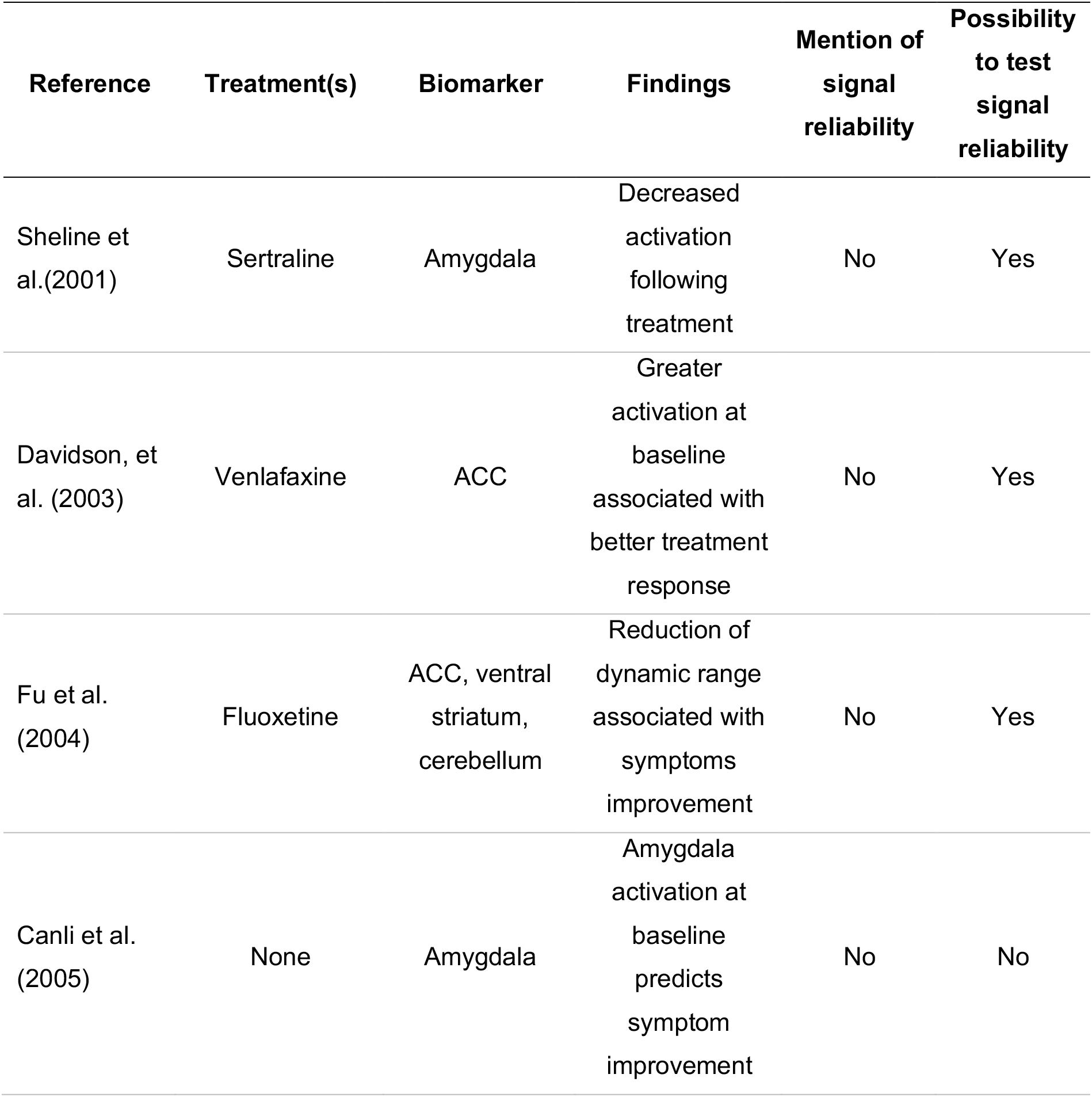
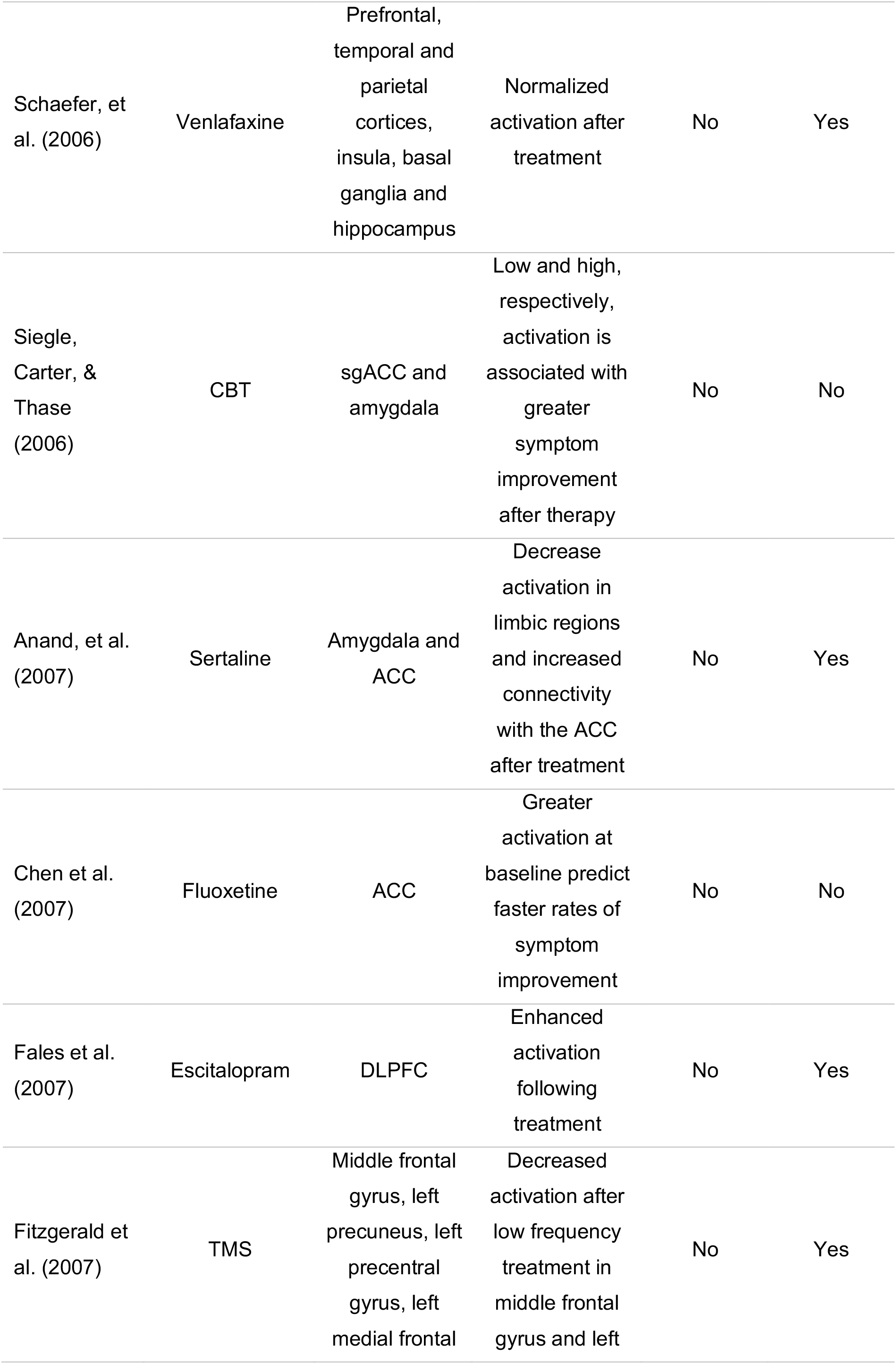
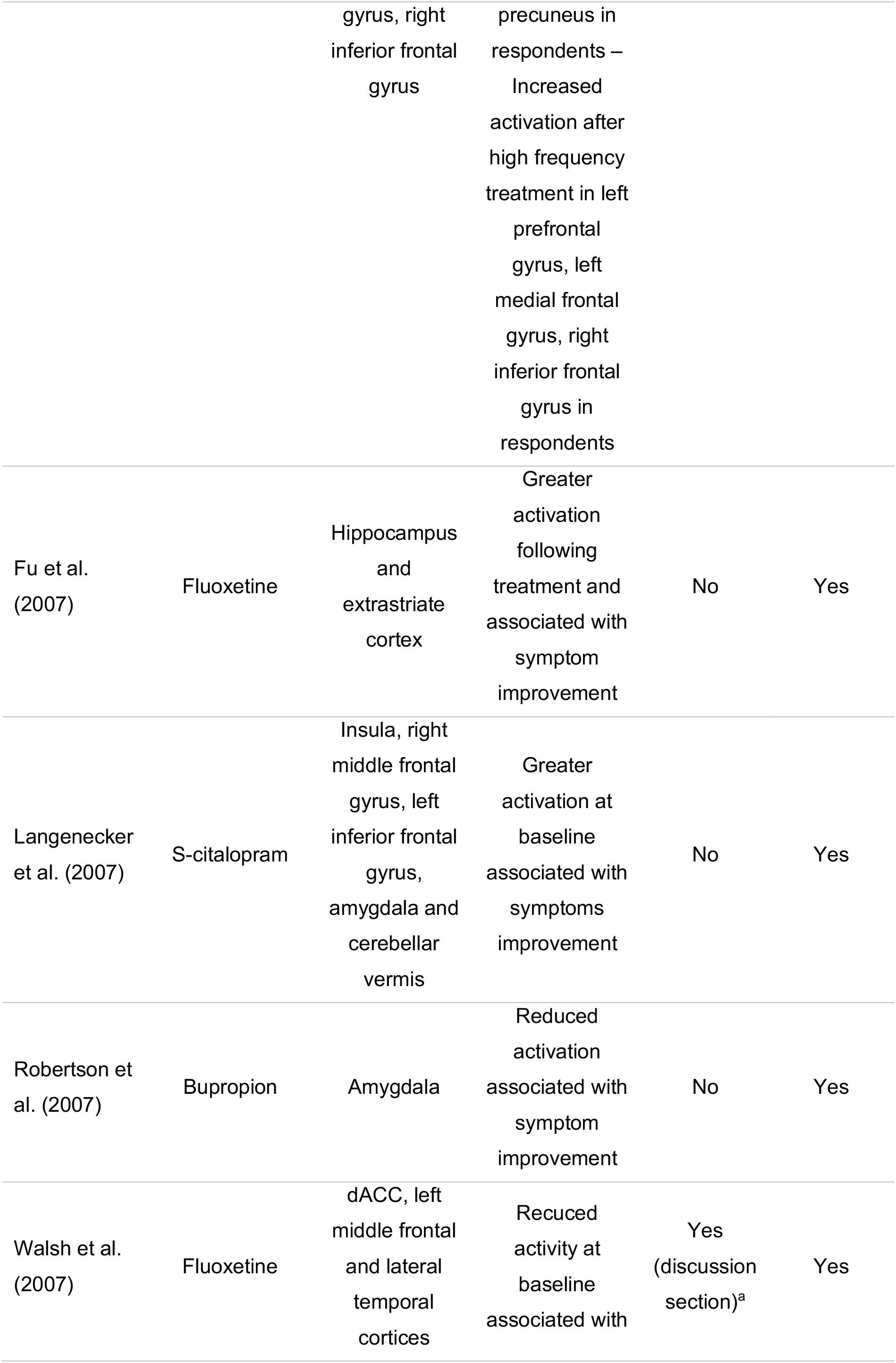
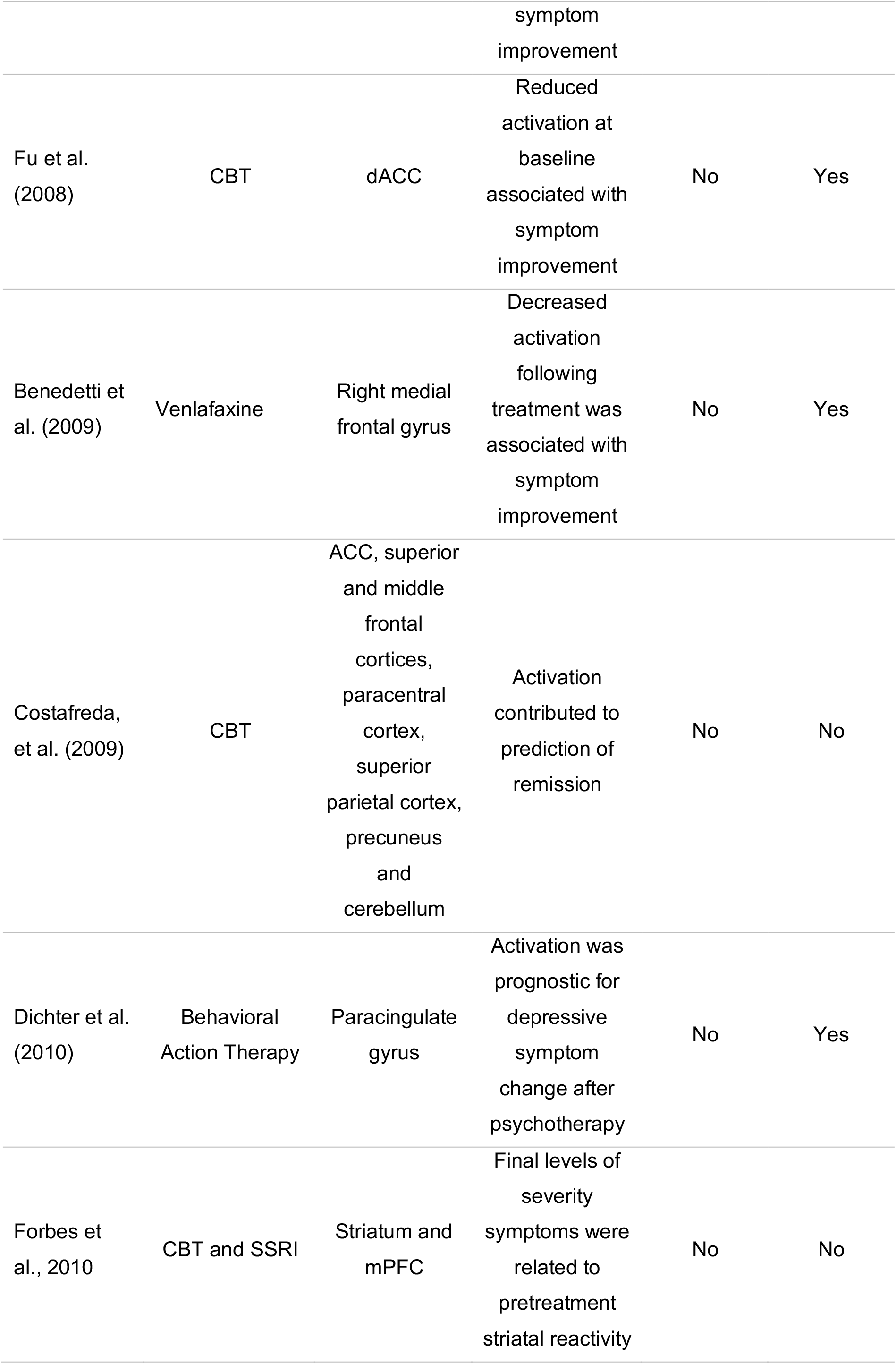
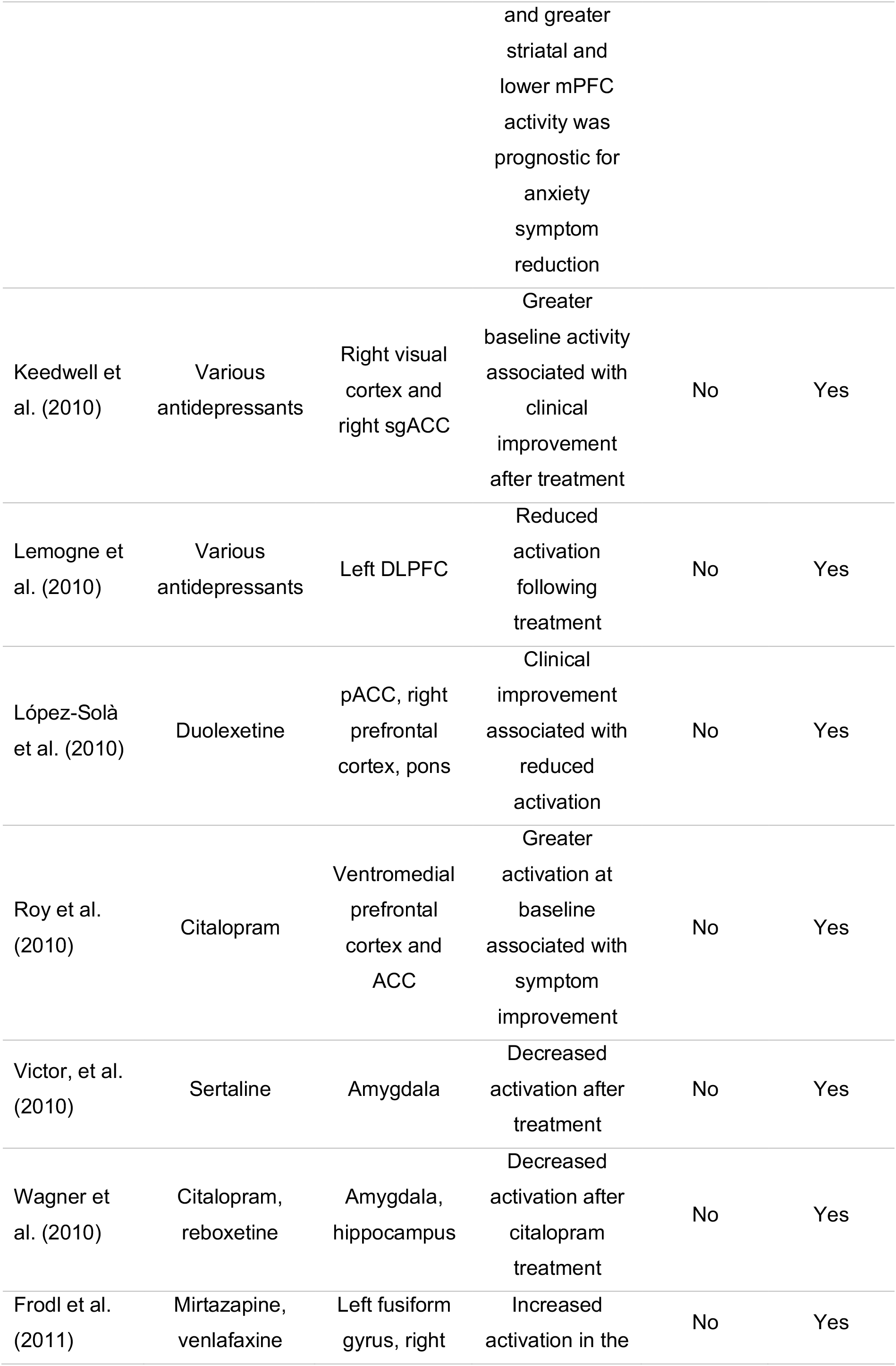
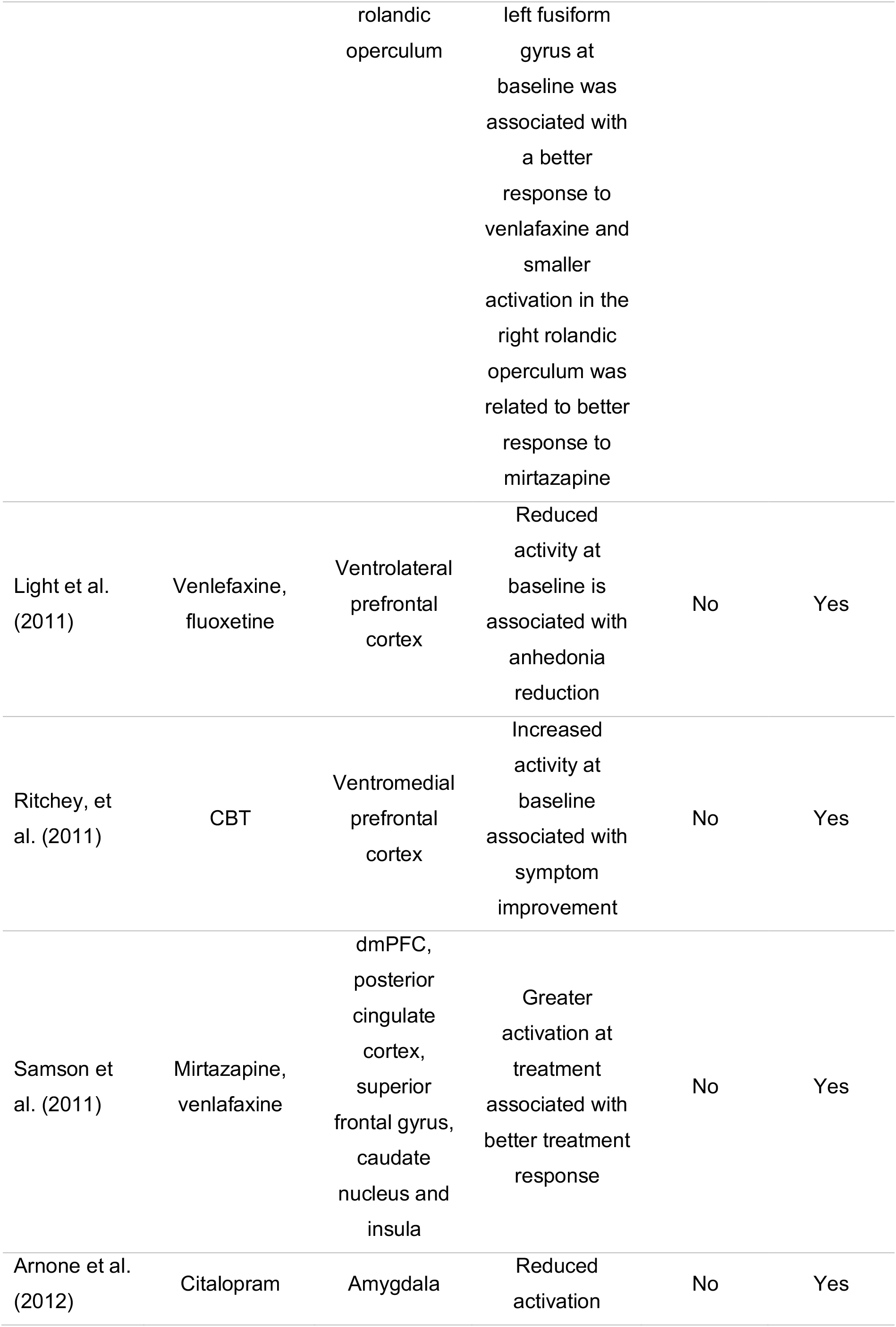
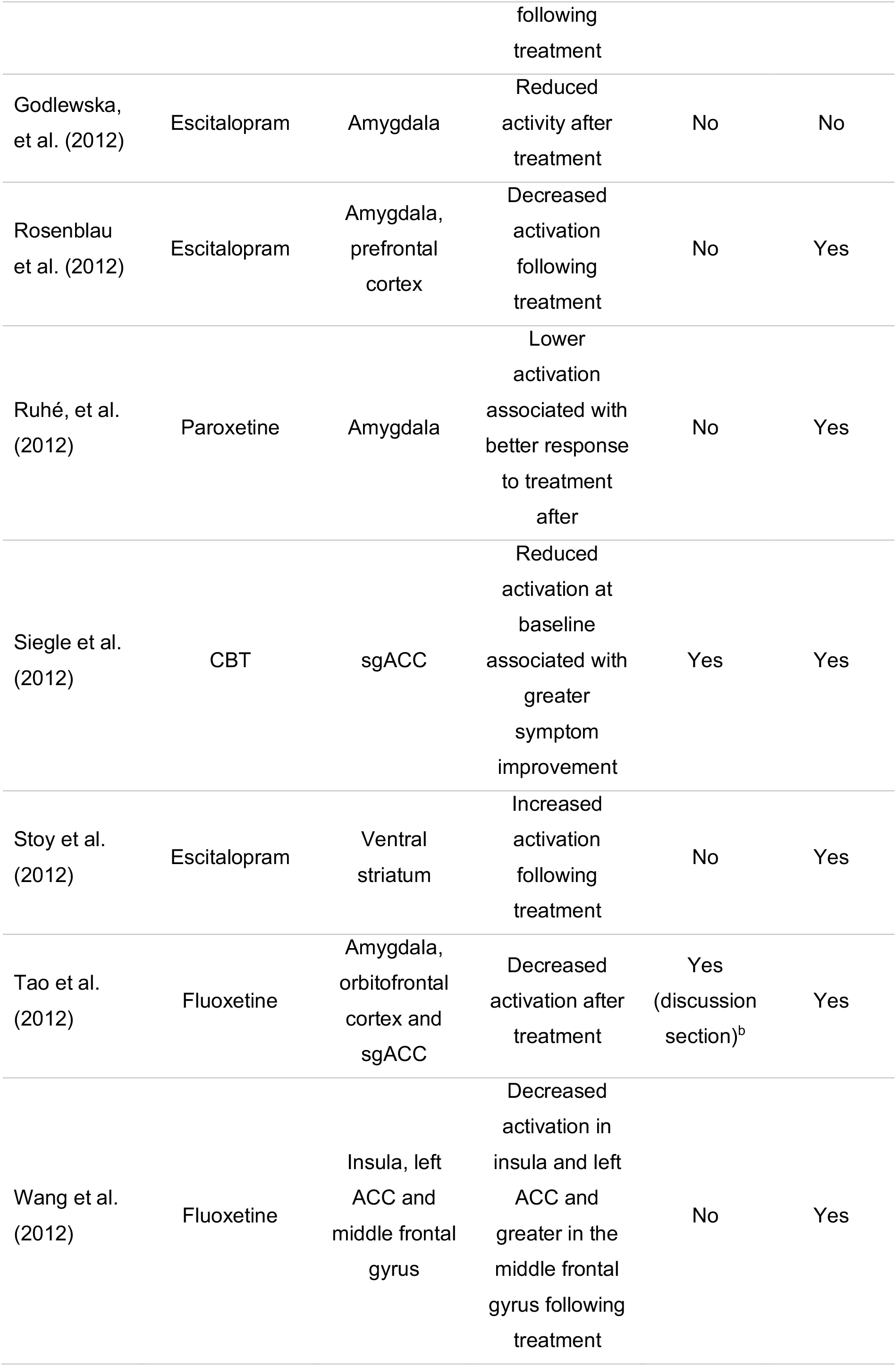
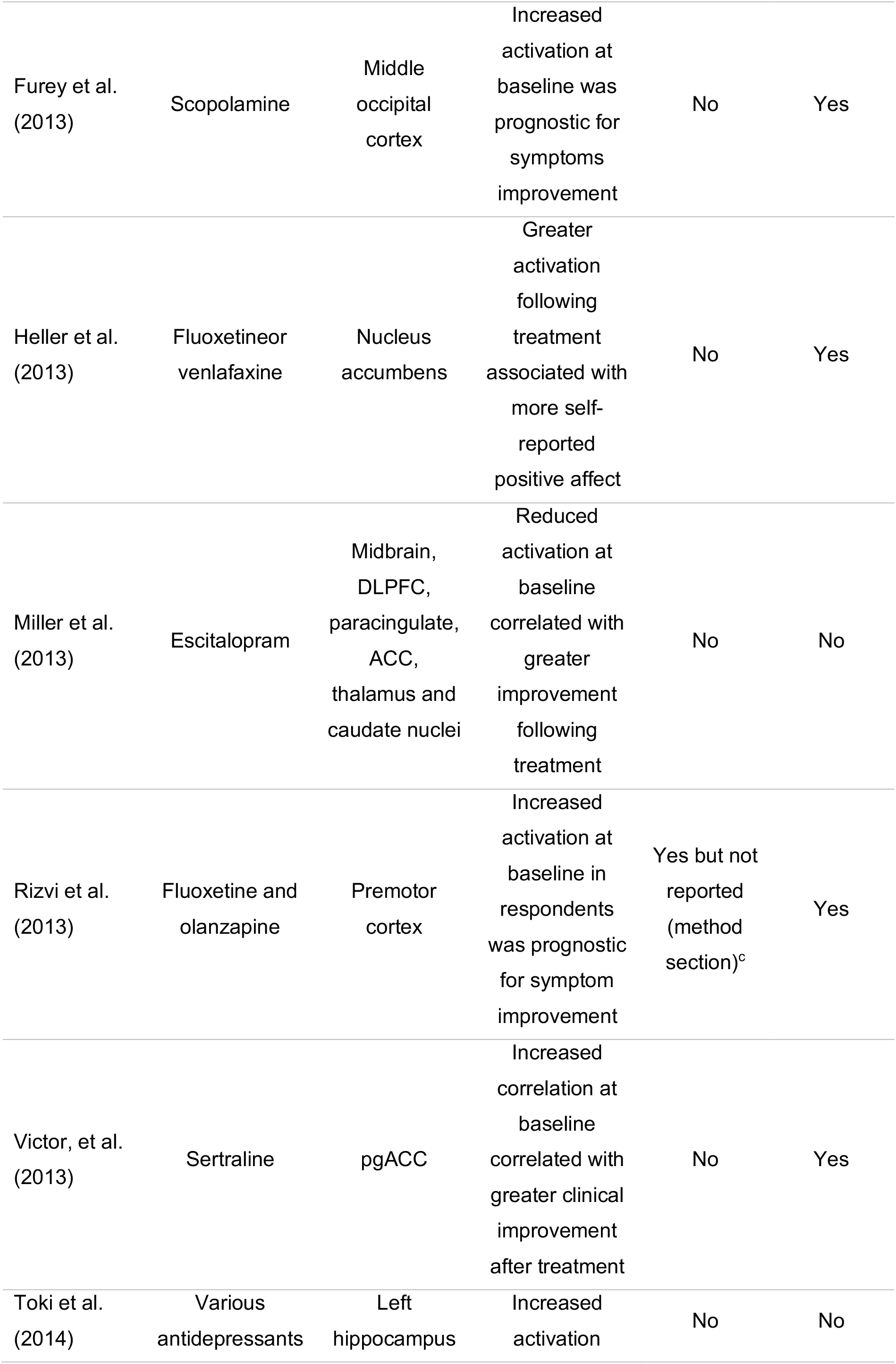
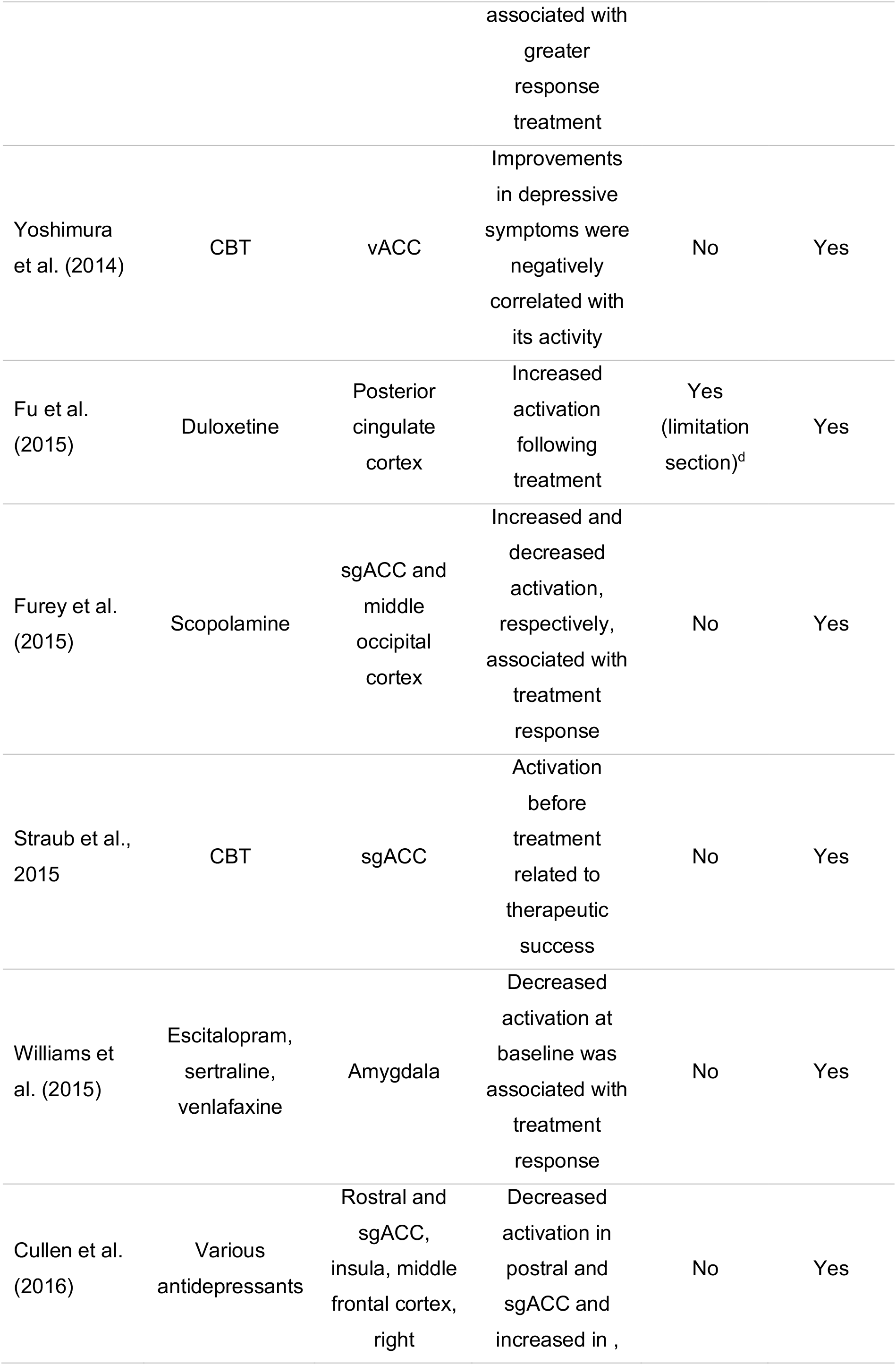
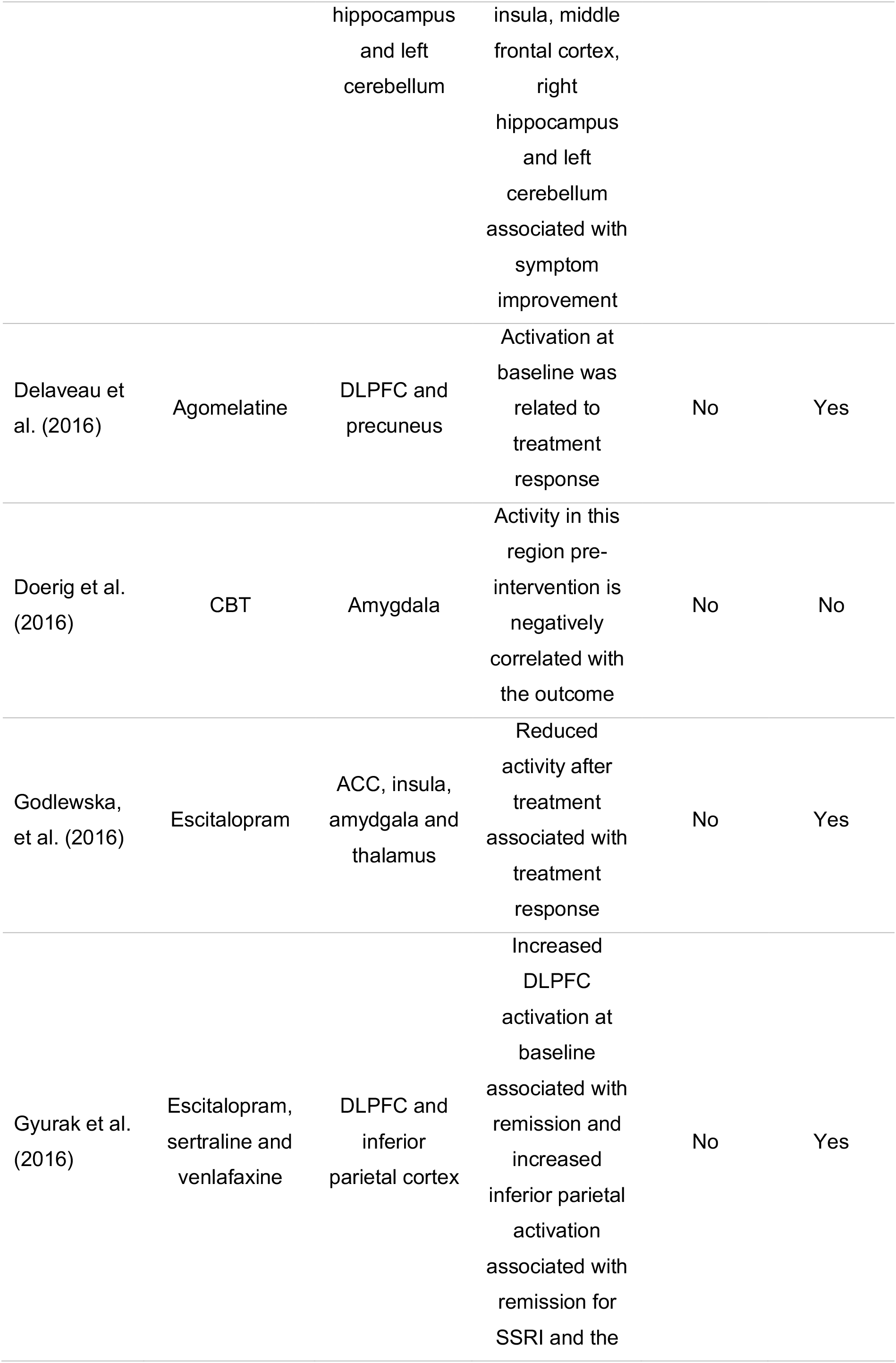
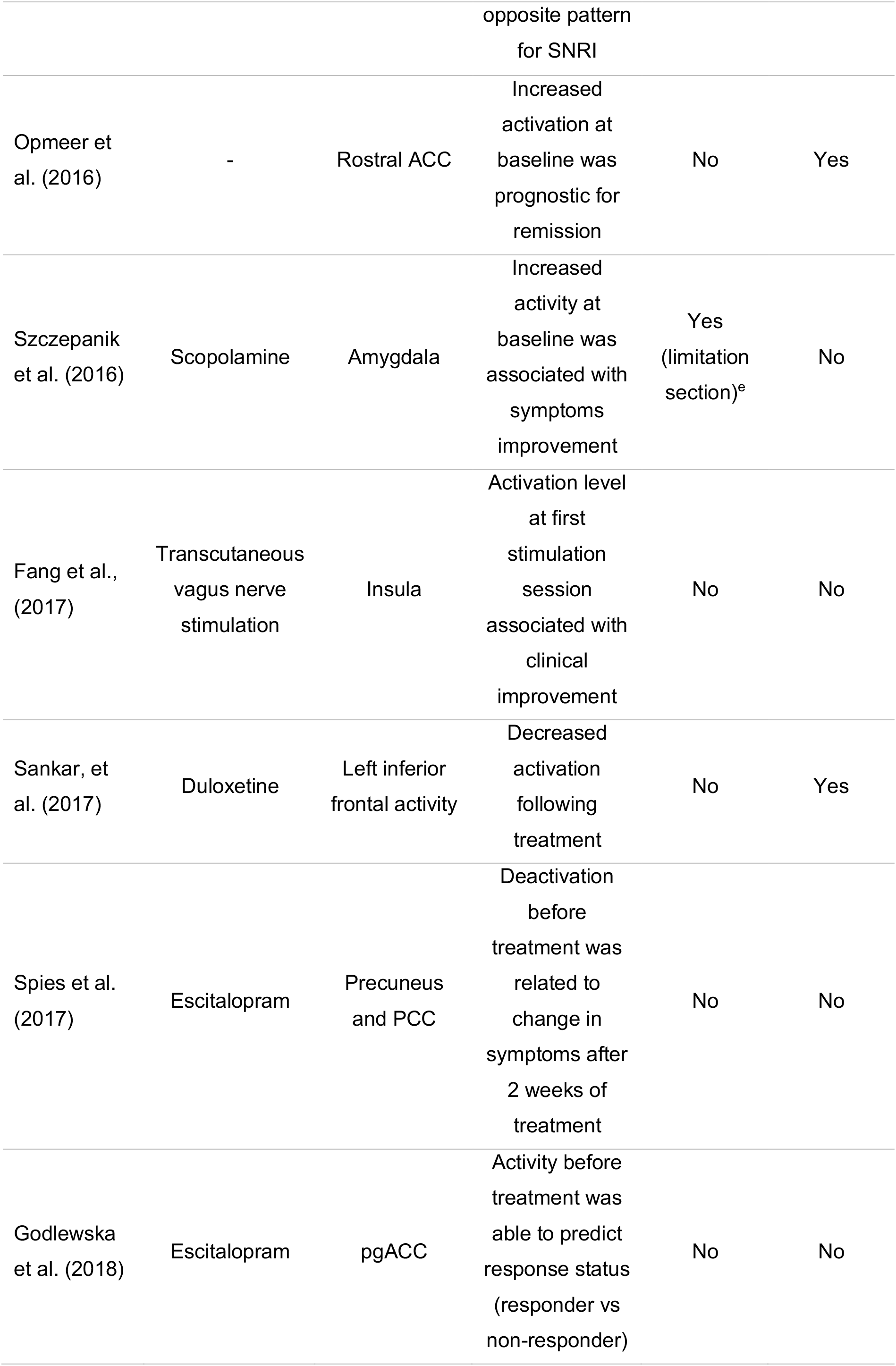
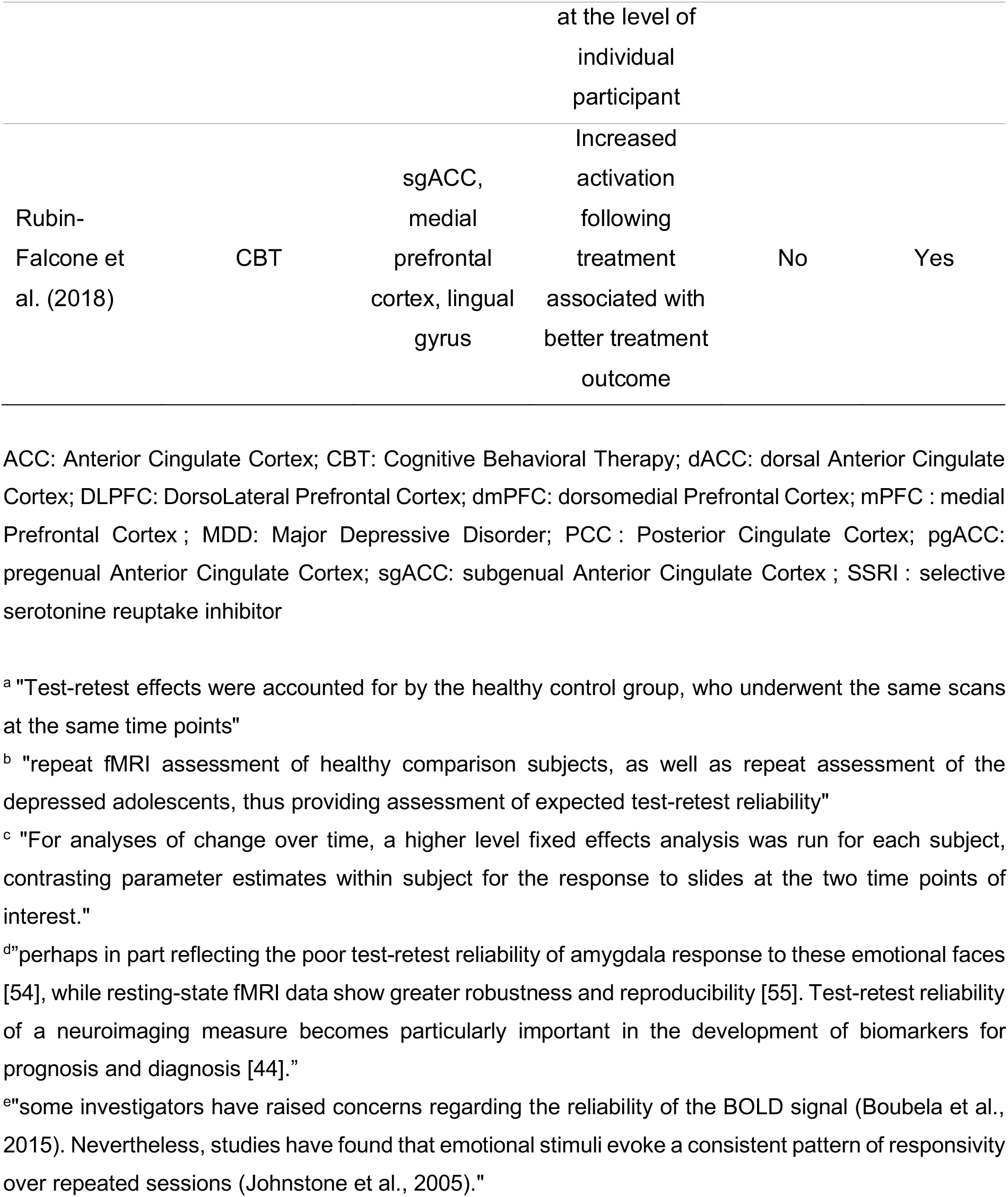
Studies Examining neuroimaging biomarkers of pharmacotherapy and psychotherapy outcomes in Major Depressive Disorder and mention of test-retest reliability of the studies

#### 1.2.2. Results

Though most of the reviewed studies could have reported test-retest reliability (i.e., participants performed two scans), most did not mention it. Seven mentioned reliability in the discussion and only one reported test-retest reliability at the subject level; Siegle et al. (2012) reported “sgACC z scores and reactivity had moderate test-retest reliability in controls undergoing testing approximately 16 weeks apart (N=27; r=0.39 [P=0.04]). All but 1 had a pretest z score less than 0.5, and all but 2 had a posttest z score less than 0.5, suggesting stability within a restricted range.” Other studies that mention reliability describe stability of group effects. For example, “Test-retest effects were accounted for by the healthy control group, who underwent the same scans at the same time points” (Walsh et al., 2007) is often reported in the discussion. This technique, while valuable, does not yield estimates of test-retest reliability at the individual subject level; the absence of a main effect of Time is evidence of the lack of a mean shift, but not of the stability of participants ranks.

### 1.3. rtfMRI-nf Studies Review

Interventions that use biological measures as real-time targets, including rtfMRI-nf also implicitly assume reliability. rtfMRI-nf trains patients to regulate the hemodynamic activity in regions of interest (Decharms, 2008) with the hope that changing a causal mechanism will result in symptom changes. rtfMRI-nf appears useful for several clinical populations, including patients with MDD (Thibault et al., 2018). Most patients can learn volitional control of hemodynamic activity in a targeted brain region (Fovet et al., 2015) which has been associated with clinical improvements (Fovet et al., 2015; Linden, 2014; Linden et al., 2012; Young et al., 2014) suggesting potential translational applications (Decharms, 2008; Ruiz et al., 2014; Thibault et al., 2018). An implicit assumption of rtfMRI-nf is that the signal measured on one day represents the same quantity measured on subsequent days, and thus performance on that metric can be trained over days. Consequently, test-retest reliability seems a strong prerequisite. Thus, as for prediction studies, we considered whether test-retest reliability is being reported in the fMRI neurofeedback literature.

#### 1.3.1. Method

A PubMed search with the key words “(neurofeedback AND fMRI) OR rt-fMRI-nf) AND (depression OR MDD OR major depressive disorder” provided 44 results. After removing articles not including rtfMRI-nf or patients suffering from MDD, we were left with 11 studies (Table 2).

**Table 2:**
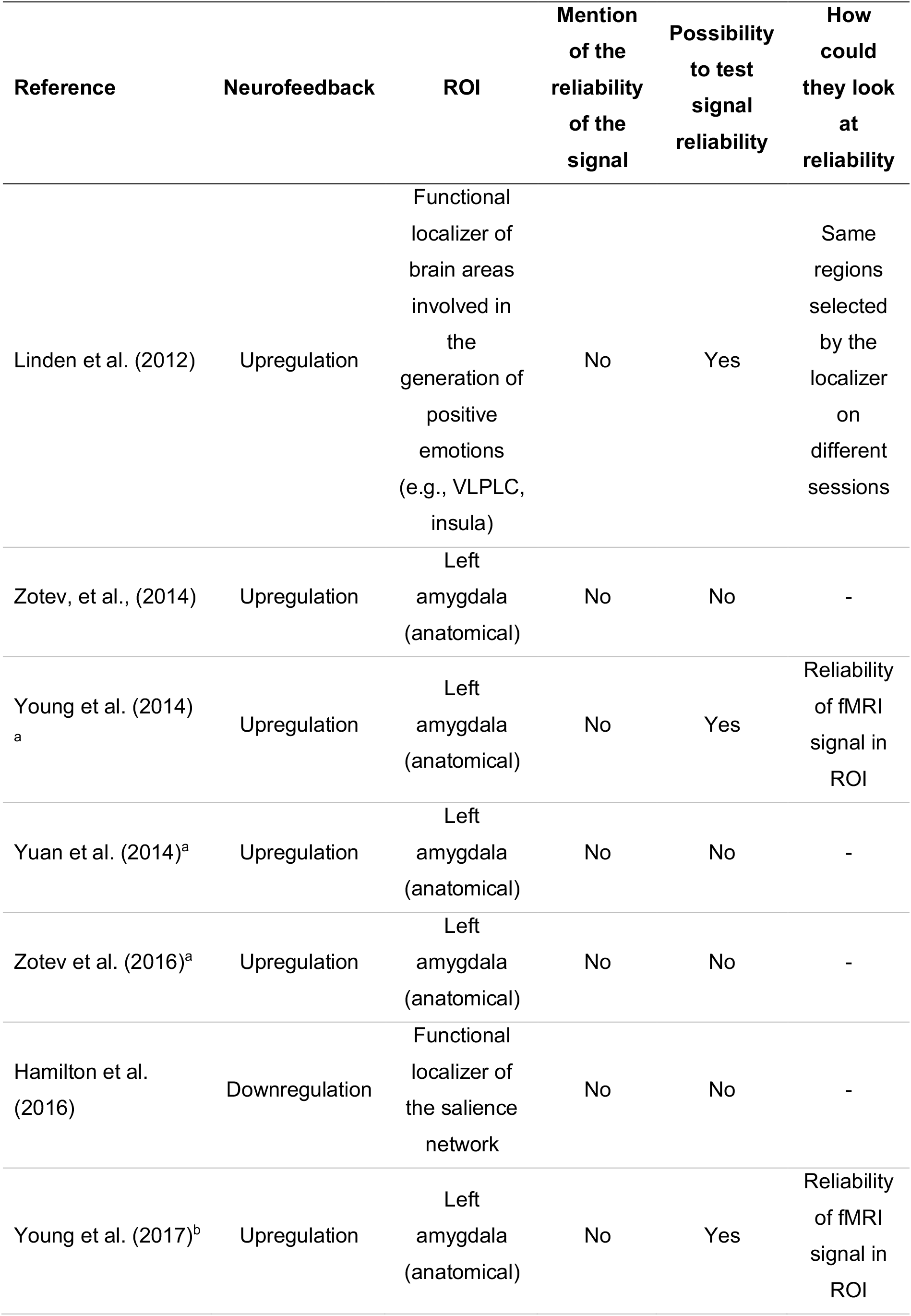
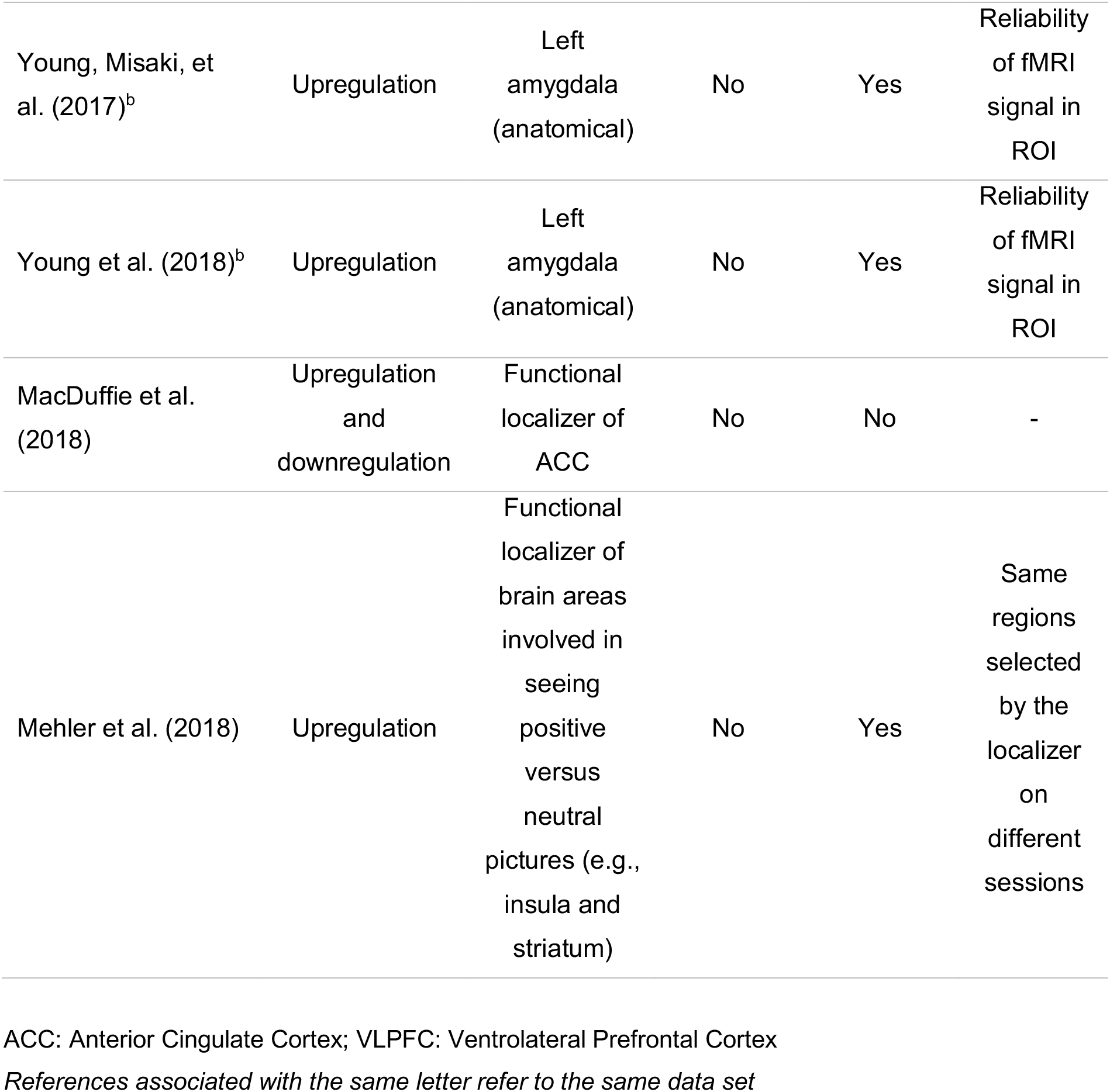
rt-fMRI-nf studies in Major Depressive Disorder and mention and possibility of test-retest reliability

#### 1.3.2. Results

None of the examined fMRI-nf studies reported on the reliability of the signal being trained (Table 2 and specific discussion of functional localizers in Box 4 in supplements).

### 1.4. Conclusions Thus Far

MDD studies using fMRI for clinical prediction or treatment rarely mention reliability, mirroring the more general fMRI literature (for meta-analysis, see Elliott et al., 2020). This lack of reporting could be due to failure to consider psychometrics important, or systematic decisions not to report observed low reliabilities. Indeed, reliability in published fMRI research in non-clinical studies, across protocols, tasks, regions of interest, psychological functions, and retest intervals have been fairly low (ICC~0.50), with most published studies reporting values between 0.33-0.66. These values are mostly below “good” reliability thresholds for psychometrically sound clinical tests (~0.6).

## 2. POTENTIAL WAYS TO OPTIMIZE TEST-RETEST RELIABILITY IN fMRI/rtfMRI-NF

To facilitate reporting of reliability in clinical studies as part of every-day neuroimaging-science, the remainder of this article is dedicated to introducing ways to report, improve, and increase clinical applicability of test-retest reliability for fMRI in clinical populations. We apply and evaluate these suggestions in two published data sets (Siegle et al., 2012; Young et al., 2017b).

There is already a strong literature on optimizing preprocessing, which can increase measurement of true signal, and thus reliability (Andersson et al., 2001; Miki et al., 2000; Oakes et al., 2005; Zhilkin and Alexander, 2004). We therefore begin by considering whether using alternate ways of indexing task-related reactivity in single-subject data with optimized preprocessing lead to improved test-retest reliability.

As each combination of task, design, scanner, preprocessing and analysis strategy has a unique value of reliability that cannot necessarily be generalized to other studies (Braver et al., 2010), it may be useful to have standardized generally applicable methods to find out which regions and analysis methods have sufficient psychometric qualities to be used as biomarkers or in which the signal is stable enough to be able to give relevant feedback of its activation.

### 2.1. Optimize indices of task-related reactivity

The first possibility we consider involves optimizing indices for task related reactivity in fMRI. This Blood Oxygen Level Dependent (BOLD) response is generally considered to be convolution of the time-course of neural activity with a physiological hemodynamic response. Mis-specification of the shape of BOLD reactivity can introduce inefficiency and noise into estimates, which decreases reliability in human (Handwerker et al., 2012; Lindquist et al., 2009; Shan et al., 2014) and animal models (Peng et al., 2019). If, for example, neural responses to task stimuli are sustained in depression rather than increased in amplitude (e.g., Mandell, Siegle, Shutt, Feldmiller, & Thase, 2014), standard indices such as the amplitude of the canonical BOLD response may not capture relevant aspects of the pathology.

Thus, we propose evaluating indices such as the average amplitude, area under the curve and timing/shape of the curve of the BOLD response in addition to its canonical amplitude. Gamma variate models, in particular, yield parameters for onset, rise and fall slopes, and magnitude of hemodynamic responses (e.g., Larson et al., 2006), which can be evaluated for reliability. Similarly, including temporal and dispersion derivatives can account for individual differences in peak response timing and small differences in HRF length, providing larger test-retest reliability values (Fournier et al., 2014).

### 2.2. Examine Regions with Voxel-Wise High Test-Retest Reliability

When considering task-related reactivity in a region of interest (ROI), it is useful to reduce voxelwise reactivity to a single or few indices which capture reactivity across the region as a whole. The same consideration applies for reliability. Caceres et. al. (2009) suggest computing the ICC in each voxel within a region of interest (ROI) and reporting the median ICC as an index of region’s test-retest reliability. This approach has been applied practically (Fournier et al., 2014; Lois et al., 2018). However, several potential biomarkers and neurofeedback targets identified in the literature, including the amygdala (Lebow and Chen, 2016; Young et al., 2014) and the sgACC (Siegle et al., 2012), consist of subregions with anatomical and functional heterogeneity (Hrybouski et al., 2016; Palomero-Gallagher et al., 2019). Their reliability may not be the same across these sub-divisions (Brabec et al., 2010; Janak and Tye, 2015; LeDoux, 2012). Therefore, it is possible that only some parts of ROIs may have adequate reliability and that the median reliability will not capture the most reliable parts of the signal. Just as questionnaires are traditionally constructed by eliminating unreliable items from an initial theoretically plausible set (Sheatsley, 1983), an index that inherits solely from the reliable voxels may increase psychometric properties of the preserved portions of regions.

Ten years ago, Bennet and Miller (2010) suggested that voxelwise reliability constitutes the most rigorous criteria of reliability since it implies that the level of activity in all voxels should remain consistent between scans. Although few studies have used this approach, we contend the available psychometric arguments weight in favor of voxel-wise computation of ICCs, restricting “reliable” ROIs to those regions in which all voxels have good or excellent reliability.

### 2.3. Optimize Models to Account for Individual and Clinical Features

Minimizing sources of non-interest that could vary between administrations increases the reliability of acquired data (Lin and Monica Way, 2014). Some fMRI noise sources such as differences in instrumentation, time of day, motion, etc. can be controlled, to some extent, via design. Tasks can be selected which have few practice effects and pre-baseline training can remove practice and strategy-development effects (Barch and Mathalon, 2011; Palmer et al., 2018). Choosing as-simple-as-possible tasks can minimize the impact of non-task cognitive processes. Standardizing instructions and training procedures helps to ensure participants understand the task before the first administration (Barch and Mathalon, 2011). Effects of other time-varying noise sources, such as thermal and physiological noise, are routinely minimized via preprocessing procedures (Krüger and Glover, 2001).

That said, if sources of variation across time, such as physiological or cognitive features, cannot be fully managed within design or processing, statistical methods for adjusting test-retest reliability estimates for them (Atri et al., 2011; Hsiao et al., 2011; Laenen et al., 2006) may be important to consider. Indeed, individual differences in state anxiety can account for amygdala activation (Calder et al., 2011) and habituation (Sladky et al., 2012), and, variation in rumination in depression is continuously associated with individual differences in amygdala, hippocampal, and prefrontal reactivity to emotional stimuli (Mandell et al., 2014; Siegle et al., 2002). Thus, true signal differences due to anxiety, mood or other symptoms between scans, especially if test-retest reliability is being evaluated in the context of possible treatment-related effects, might account for apparently unreliable neural responses, particularly to emotional stimuli. Thus, it may be useful to account for individuals’ differences that could change across time statistically in estimating reliability, e.g., via the inclusion of clinical covariates.

### 2.4. Examine Reliability Within Relevant Tasks and Clinical Populations

Estimating reliability in healthy participants or symptomatic individuals who do not receive intervention may help separate effects of symptom change from practice effects. Yet, these approaches can introduce other confounds (e.g., if a task is reliable in patients but not controls or non-treatment seeking symptomatic individuals). The majority of studies have examined reliability of fMRI data in homogenous samples of healthy, often young, university students (Bennett and Miller, 2010; Lois et al., 2018). Studies reviewed in Table 1 that discuss reliability in MDD generally restrict their discussion to whether there was a main effect of Time in healthy controls. Generally, BOLD response variability is greater in forms of between-subject responses than within (Aguirre et al., 1998). A limitation of ICC is that simultaneous inclusion of within and between subject variability causes estimators to be affected by sample composition. As groups might differ in the degree to which regional signals are reliable between measurements (Fournier et al., 2014), and because ICCs are proportional to between subject variability, heterogeneous samples can produce different ICCs even with the same degree of within-subject reliability of test-retest values. Using only healthy control participants may underrepresent true variability or over represent measurement errors in the population of interest, yielding inaccurate reliability estimates. Similarly, non-treatment seeking patients differ from treatment seeking patients on many variables that could affect test-retest reliability, such as symptomatology and comorbidity (Galbaud Du Fort et al., 1999).

Thus, testing reliability in the population of interest may provide more accurate estimates. We therefore recommend the use of representative samples to create a voxel-wise, population- and task-specific map of test-retest reliability. For example, if a task is to be used to distinguish symptomatic from healthy individuals, this method should be applied to a mixed population of healthy and symptomatic participants prior to the clinical application of the task. If the purpose is to distinguish respondents and non-respondents to a treatment, we recommend assessing reliability among treatment-seeking patients.

## 3. EVALUATION OF SUGGESTED OPTIMIZATIONS IN A PROGNOSTIC NEUROIMAGING TREATMENT OUTCOME DATASET

We have described several approaches that could be useful when examining and seeking to improve test-retest reliability in service of clinical translation including R1) optimizing BOLD signal parameterization, R2) using regions or voxels with stronger psychometric properties, R3) accounting for within-individual changes and R4) studying relevant tasks and populations for the intended application. In this section we demonstrate feasibility of these approaches and examine whether they are useful when applied to a published clinical fMRI dataset (Siegle et al., 2012). Our code for these analyses is freely available from https://github.com/PICANlab/Reliability_toolbox in the folder named “activation_task_reliability”.

### 3.1. Method

The sample consisted of participants described in Siegle et al. (2012) augmented by the addition of 8 patients who completed the same protocol after that paper was submitted, yielding 57 patients with major depressive disorder (MDD), and 35 healthy control participants (see supplement for details of this dataset and its relationship to Siegle et al 2012). Briefly, participants with MDD completed a slow event-related task during 3T fMRI in which they labeled the valence of emotional words (here, as in the published dataset, we analyzed only nominally negative words) before and after 12-16 weeks of Cognitive Therapy.

We computed reliability estimates within 4 ROIs which the literature suggests may function as biomarkers for treatment response including the amygdala (Arnone et al., 2012; Godlewska et al., 2012; Sheline et al., 2001), dorsolateral prefrontal cortex (DLPFC, Koenigs and Grafman, 2009), rostral anterior cingulate cortex (rACC, Hunter et al., 2013) and subgenual cingulate cortex (sgACC, Siegle et al., 2012b; Straub et al., 2015; Taylor et al., 2018) (our region-wise definitions are included in Box 1 in Supplement).

#### 3.1.1. Optimize the BOLD Signal

The BOLD response to negative words was modeled within participants using 4 different methods including 1) amplitude of a canonically shaped BOLD signal (using AFNI’s 3dDeconvolve with a narrow tent function (‘BLOCK5(1,1)’, Cox, 1996), 2) Area under the curve (via multiple regression of a delta function across 8 TRs using 3dDeconvolve, i.e. computed with Finite Impulse Response/FIR basis, with sum of betas as the parameter retained); 3) Peak amplitude from the same regressions as #2, and 4) a gamma variate model with parameters for onset-delay, rise-decay rate, and height. Voxelwise outliers outside the Tukey hinges were windsorized across participants and ICCs (3,1) were computed (Shrout and Fleiss, 1979) within individuals for each modeling method using custom Matlab code. While ICC(2,1) allows generalizing results obtained from different scanners, we chose to use ICC(3,1) to be able to compare with most of the literature, given that it is the most widely used ICC. This approach also allowed us to examine the importance of including scanner as a covariate in 3.1.3.

#### 3.1.2. Compute Voxelwise Reliability

To measure the benefit of identifying reliable voxels, we calculated the mean, median and standard deviation of the ICCs in each of the ROIs for each modeling method and each group.

#### 3.1.3. Include Clinical and Design Related Measures

We examined whether indices of reliability increased when clinical and design-related measures were included. As the ICC does not easily allow inclusion of covariates, we used semi partial correlations within the context of multiple regressions with and without covariates to assess changes in reliability, where covariates were pre and post clinical measures, as:

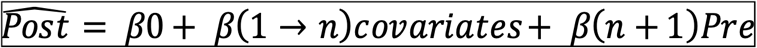

This model accounts for the potential that participants who show little change in symptoms may have better test-retest reliability. Modelling these clinical effects at the group level should make it possible to identify variance unique to test-retest reliability.

We included indices of pre- and post-treatment depressive symptomatology (Beck Depression Inventory; BDI, Beck et al., 1996), state and trait anxiety (Spielberger, 1983), rumination (Nolen-Hoeksema et al., 1993), and sleepiness (Johns, 1991) administered on the scan day, the scanner on which data were acquired, and participant’s group when patients and controls were considered in one sample, coded as dummy variables, as covariates. Missing data were imputed via regression from the other administered measures also used as covariates.

A primary question was whether any of the proposed techniques described above, including different BOLD models, accounting for voxelwise variability, and the use of covariates, would differentially affect reliability estimates (i.e., semi-partial correlations). As such, after computing reliability estimates at each voxel, we rank ordered them across all permutations of BOLD estimate parameters (6 parameters) and the use or non-use of covariates (2 conditions) at each voxel per ROI, yielding 12 x #-voxels rankings per ROI. Following a Kolgomorov-Smirnov test justifying the need to use non-parametric tests, we report a Kruskal-Wallis test to determine whether the rankings differed across models in each ROI. If they did, as a simple effects test, we generated confidence intervals around the mean of rankings for each of the 12 conditions via a one-way ANOVA (via Matlab’s multcompare function). Non-overlapping confidence intervals are interpretable as significant differences between one condition and any other. To display them we generated figures showing the mean of rankings for each condition, which will be numbers on the order of 1 to 12 x # voxels, with higher means representing being at the top of the rankings across many voxels within the ROI.

#### 3.1.4. Use Clinically Representative Samples

All analyses were conducted on the whole sample (controls and patients) to establish likely reliability of tests that could be used to discriminate groups, and on patients only, to establish likely reliability of clinical prognostic and change indicators. We considered multiple reliability effect size thresholds which might be used in other studies (0.4 and 0.6 for fair and good reliability and 0.7, and 0.75 for traditional labels of the data as “reliable” and clinically meaningful).

#### 3.1.5. Type 1 error control

As 1) each of the hypotheses and regions examined for this manuscript was considered a different family of tests and 2) we want our results to generalize to reliability as it is reported in the confirmatory biomarker and neurofeedback literatures where only one region is generally examined, consistent with the literature on test-retest reliability in neuroimaging, type I error was not controlled across regions and hypotheses for ROI-wise statistics. For simple-effects tests of differences in rankings across conditions, we controlled for the number of conditions with a Bonferroni test. For voxelwise statistics we subjected all voxelwise residual maps to empirical cluster thresholding (AFNI’s 3dFWHMx and 3dClustSim, acf model with small-volume corrections for examined regions) using a p threshold (-pthr) based on each considered effect size threshold (see in supplementary materials, table S3 for more details).

### 3.2. Results and discussion

#### 3.2.1. Optimizing the BOLD signal

ICC’s were uniformly low (<.3) for all BOLD parameterizations when entire ROIs were considered (Table 4). Kruskal Wallis tests did suggest differential reliability across our parameterizations (Table 5a). This held when the two outlying uniformly low reliability parameterizations (rise decay with and without covariates) were removed from consideration (Table 5b). Yet, there were non-overlapping confidence intervals among counts of rank orderings of parameterizations for voxelwise tests, suggesting that at least for some subsets of regions, some parameterizations were superior (Supplement Figure S1, Table S1). For example, in the full sample, for the amygdala, amplitude without covariates was superior to other parameters. Over all ROIs, the most reliable parameters were amplitude, canonical amplitude, and height (Figure 1 shows voxelwise variation within a Priori ROIs for the height parameter) for the whole sample and amplitude, area under the curve, and height for only patients (Figure S1 and Table S1). However, looking at ROIs and samples independently, the parameter offering the highest levels of reliability varied.

**Figure 1:**
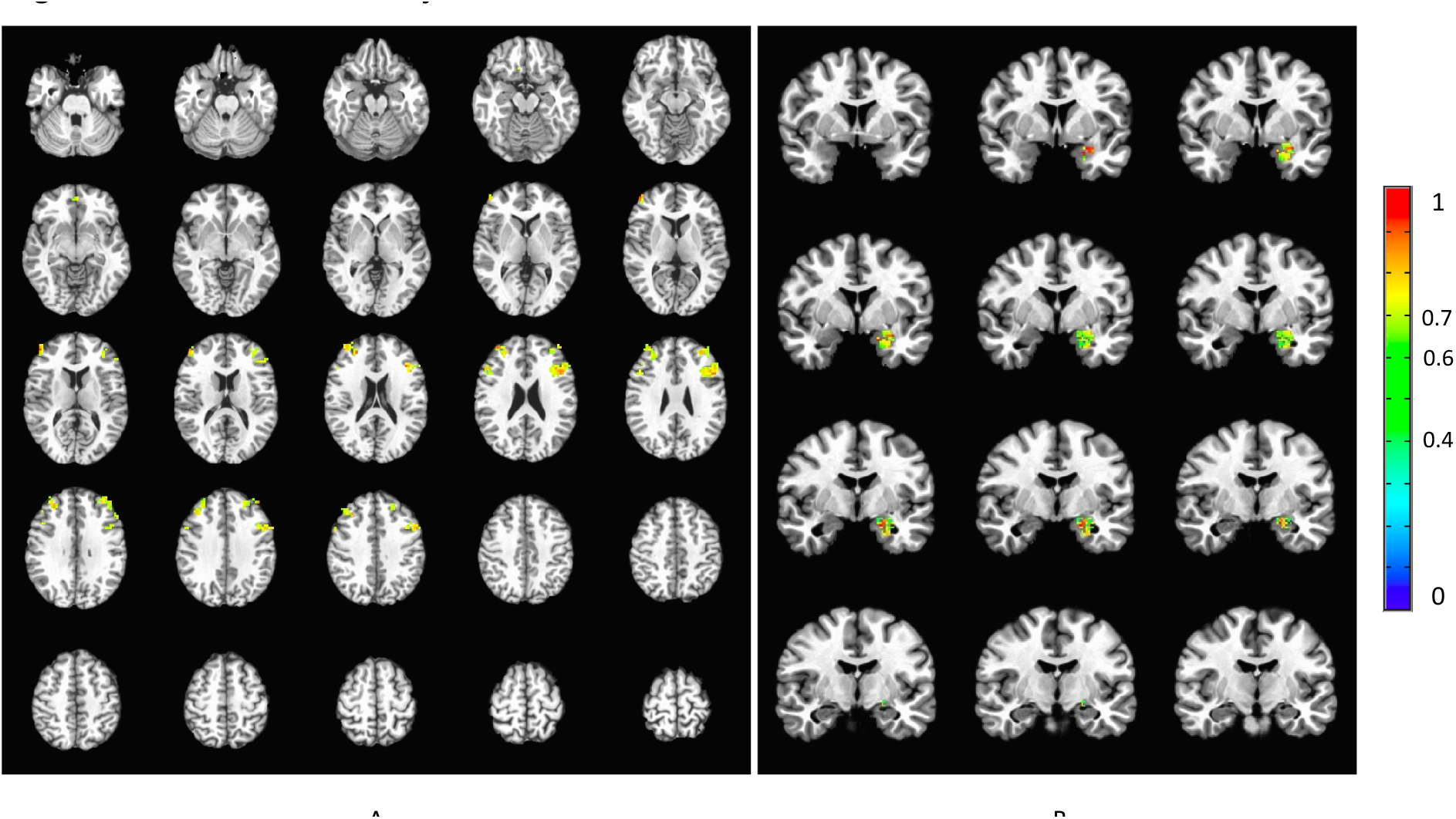
Test-retest reliability in ROIs estimated with voxel wise ICCs using height parameter, a threshold of ICC>0.4 and cluster correction applied for this threshold in A. Siegle et al. (2012) dataset of patients and B. Young et al. (2017) data set of the transfer run in the experimental group (signal with training) preprocessed with the TBV style pipeline.

#### 3.2.2. Voxelwise reliability

In the whole sample, moderate reliability (ICC>.4) in clusters large enough to infer significance was observed in the DLPFC using the canonical amplitude model and in the amygdala using amplitude (Table 3). “Good” (ICC>.6) reliability was reached in clusters large enough to infer significance when only the patients were considered, using amplitude and height in the DLPFC. These levels of voxelwise test-retest reliability were higher than using the median or mean value of ICCs within whole ROIs (Table 4). Levels of generally accepted reliability for clinical measures (ICC>.7) were not observed in clusters large enough to report.

**Table 3:**
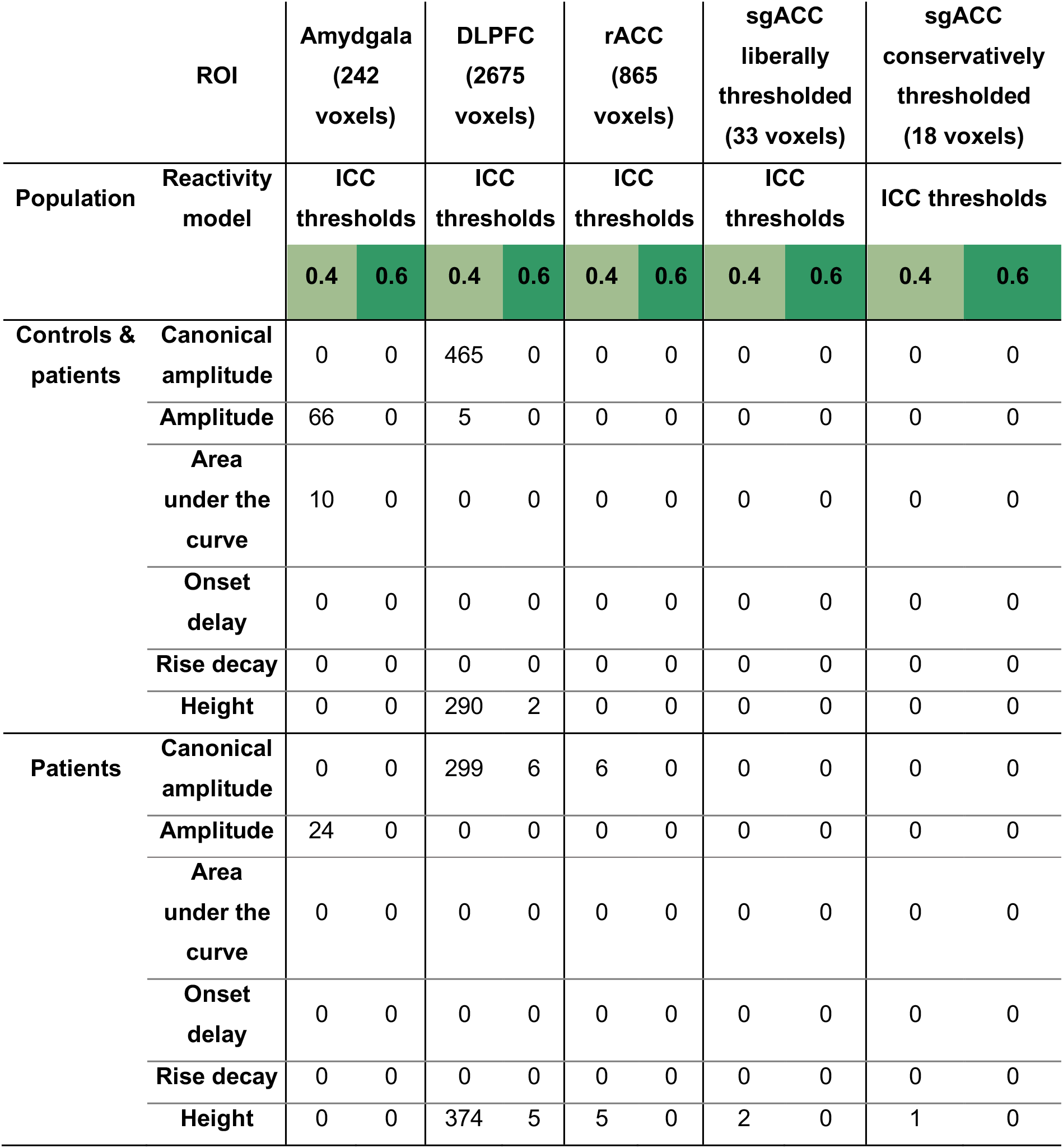
Table of number of voxels reaching different reliability thresholds for each sample, first level parameter, and ROI with cluster correction applied.

**Table 4:**
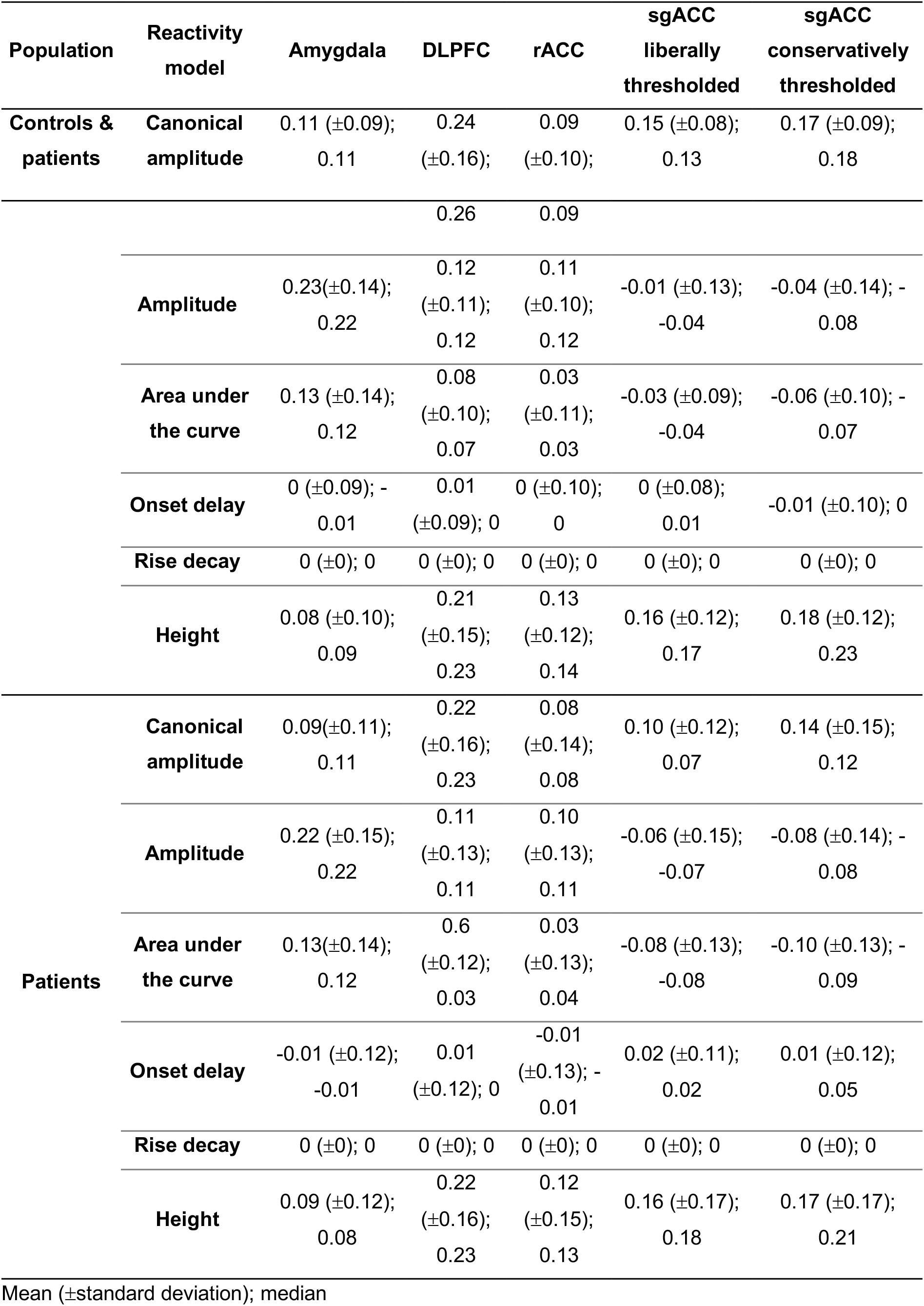
Table of mean, standard deviation and median values of ICCs for each sample, reactivity model, and ROI.

**Table 5a:** Table of Kruskal Wallis tests’ output for each sample, reactivity model with and without covariates, and ROI with Bonferroni correction applied.

| Population | Amygdala | DLPFC | rACC | sgACC<br>liberally<br>thresholded | sgACC<br>conservatively<br>thresholded |
| --- | --- | --- | --- | --- | --- |
| Controls &<br>patients | H(11)=1414.67,<br>p<0.001 | H(11)=12717.07,<br>p<0.001 | H(11)=4794.14,<br>p<0.001 | H(11)=206.47,<br>p<0.001 | H(11)=118.32,<br>p<0.001 |
| Patients | H(11)=1233.13,<br>p<0.001 | H(11)=10371.75,<br>p<0.001 | H(11)=4477.55,<br>p<0.001 | H(11)=240.89,<br>p<0.001 | H(11)=136.93,<br>p<0.001 |

**Note: Applying Bonferroni correction for 6 reactivity models with and without covariates ( $p < 0.05/12 = 0.004$ ).**

**Table 5b:** Table of Kruskal Wallis tests’ output for each sample, reactivity model with and without covariates, and ROI with Bonferroni correction applied, without rise decay.

| Population | Amygdala | DLPFC | rACC | sgACC liberally thresholded | sgACC conservatively thresholded |
| --- | --- | --- | --- | --- | --- |
| Controls & patients | H(9)=285.88,<br>p<0.001 | H(9)=4876.99,<br>p<0.001 | H(9)=644.19,<br>p<0.001 | H(9)=58.90,<br>p<0.001 | H(9)=40.55,<br>p<0.001 |
| Patients | H(9)=25.68,<br>p=0.002 | H(9)=1588.15,<br>p<0.001 | H(9)=190.42,<br>p<0.001 | H(9)=108.21,<br>p<0.001 | H(9)=67.20,<br>p<0.001 |

**Note: Applying Bonferroni correction for 6 reactivity models with and without covariates ( $p < 0.05/10 = 0.005$ ).**

#### 3.2.3. Clinical and Design Related Measures

The addition of covariates never resulted in significantly higher average ranks for semi partial correlations in any ROI, in the whole sample or just the patients (Figure S1). In other words, adding covariates did not improve the reliability, and in some instances made it worse.

## 4. EVALUATION OF SUGGESTED OPTIMIZATIONS IN AN EMPIRICAL NEUROFEEDBACK DATASET

To further support the feasibility of applying these recommendations and to evaluate the consistency of their performance in a second dataset, we consider a published fMRI neurofeedback dataset (Young, Siegle, et al., 2017, code available from https://github.com/PICANlab/Reliability_toolbox in the folder named “rtfMRI-nf_reliability”).

### 4.1. Method

This dataset constituted 18 patients in the experimental group who received amygdala neurofeedback and 16 patients in the control group who received parietal neurofeedback. Briefly, participants completed two training scans on different days within 2 weeks, each including a “baseline” and “transfer” runs during which no feedback was presented. The analyzed task was a 40-second per block design during which participants alternately rested, worked to upregulate a target region during recall of positive memories, and did a distraction (counting) task (see supplement Box 5 for details of this dataset). Here, we focus on a) the baseline data on the two training days in control-feedback participants during recall of positive autobiographical memories prior to neurofeedback training. As their amygdala signal did not change over the course of the study at the group level (Young et al., 2017b), this allows us to examine test-retest reliability of the left amygdala signal without the influence of neurofeedback. b) the left amygdala signal during the two transfer runs in the experimental group, as this represents the effect of neurofeedback training. Activity during the two post-training transfer runs did not differ at the group level, allowing us to examine the test-retest reliability of the amygdala signal after neurofeedback training. Because this dataset only included patients with MDD, only the first 3 principles (i.e., optimization of the BOLD signal, computation of voxelwise reliability, and inclusion of clinical and design related measures) are evaluated in this dataset.

#### Feedback signal

To analyze the feedback signal averaged over the left amygdala we used the output of the script used in Young, Siegle, et al. (2017) that allowed computation of the feedback signal in real-time before considering the voxel-wise signal.

#### Voxel-wise

As rtfMRI-nf involves real-time preprocessing of the data, we sought to examine whether this kind of preprocessing could affect the test-retest reliability of the signal. We therefore performed data preprocessing emulating the real-time data processing performed by the commercially available neurofeedback software Turbo BrainVoyager (BrainVoyager, The Netherlands; henceforth “TBV style”) and a more classic contemporary post-hoc preprocessing stream (here referred to as “standard preprocessing”). Both streams were implemented using AFNI.

##### TBV style preprocessing

Turbo BrainVoyager performs the following functions in real-time: 3D motion correction, spatial smoothing, and drift removal via the design matrix. We used AFNI to approximate these steps. After spatially transforming the anatomical then functionals to the International Consortium for Brain Mapping 152 template, we then rescaled them to conform to the Talairach atlas dimensions and then performed motion correction to the first image, spatial smoothing 4mm FWHM smoothing kernel and fourth order detrend for drift removal.

##### Standard preprocessing

MRI pre-processing included despiking, volume registration and slice timing correction for all EPI volumes in a given exam. After applying an intensity uniformity correction on the anatomical, the anatomical was spatially transformed to the International Consortium for Brain Mapping 152 template and rescaled to conform to the Talairach atlas dimensions. Then, the fMRI data for each run were warped nonlinearly and the same spatial transformations were applied. The fMRI run was spatially smoothed within the grey matter mask using a Gaussian kernel with full width at half maximum (FWHM) of 4 mm. A first standard GLM analysis was then applied separately for each of the fMRI runs. The following regressors were included in the GLM model: six motion parameters and their derivatives as nuisance covariates to take into account possible artifacts caused by head motion, white matter and cerebrospinal fluid signals, and five polynomial terms for modeling drift.

#### 4.1.1. Optimize the BOLD Signal

##### 4.1.1.1. Amygdala signal

From each participant’s real-time left amygdala signal we calculated an “amygdala signal” for each positive recall block minus the mean of the preceding rest block from the output of previously used scripts for real-time preprocessing (Young et al., 2017b), and recreated the feedback signal by taking the amount of activation at every TR during the experimental condition minus the mean activation in the previous rest condition, on the baseline run of control participants at visits 1 and 2 (signal without training) and on the transfer run of experimental participants at visits 1 and 2 (signal with training), independently. We then averaged the time course of the feedback signal over all happy blocks. We summarized the activation for each participant for each visit by either a mean of the amygdala signal or by fitting the time course with a gamma variate model with parameters for onset-delay, rise-decay-rate, and height (see Methodological choice to fit gamma variates in Supplement Box 2 for more information of this methodological choice).

ICC(3,1) estimates were computed (Shrout and Fleiss, 1979) independently on the estimates of the feedback signal with and without training.

##### 4.1.1.2. Voxelwise signal

The same reactivity models as in the treatment outcome dataset were applied (see part 3.2.1.1) to data preprocessed with both types of preprocessing but adapted to this design (AFNI tent parameters to accommodate 40 s blocks as BLOCK(40,1), and area under the curve across entire blocks).

#### 4.1.2. Compute Voxelwise Reliability

As in the treatment outcome data set, to measure the benefit of identifying reliable voxels, we calculated the mean, median and standard deviation of the ICCs in the left amygdala for each model, group, and additionally for both preprocessing pipelines.

#### 4.1.3. Include Clinical and Design Related Measures

As in the treatment outcome data set, semi partial correlations were computed with and without covariates. We included indices of depressive symptomatology (Beck Depression Inventory; BDI, Beck et al., 1996), state and trait anxiety (Spielberger, 1983), sleepiness and drowsiness administered on the scan day, and the scanner on which data were acquired coded as dummy variables, as covariates. There was no missing data. We then compared the semi-partial correlations across all models of individual responses with and without covariates for each group and preprocessing pipeline as in section 3.1.3, to understand which models offered adequate test-retest reliability and whether there were differences between them.

#### 4.1.4. Type 1 error control

As discussed in section 3.1.5, cluster correction was applied on voxelwise statistics (further details in supplement table S4).

### 4.2. Results and discussion

#### 4.2.1. Optimizing the BOLD signal

##### 4.2.1.1. Amygdala signal

The mean amygdala signals with and without training showed poor reliability (ICCs<0.1). When the amygdala signal within the left amygdala was fit using a gamma variate function, the onset-delay and height parameters showed fair reliability for the signal without training (ICC=0.54 and ICC=0.47, respectively), with all other models, including those with training, showing minimal reliability (ICC<.1). Therefore, it appears that the shape of the signal without training is consistent across sessions and that the signal in the left amygdala is more reliable when unchanged by training, which is consistent with the assumption that training is changing the signal over time.

##### 4.2.1.2. Voxel-wise signal

Kruskal Wallis tests suggested there were differences between the parameters in reliability (Tables 6a and 6b). In particular, reliability for the height parameter (as well as amplitude for the signal without training) was higher than for other parameters (Figure S1). The height parameter also yielded a large enough cluster to infer significance for “excellent” (ICC>.7) reliability in both samples (Table 7, Figure 1 for illustration).

The use of the standard preprocessing stream had non-significantly-different reliabilities from the stream emulating the real-time preprocessing run by Turbo BrainVoyager over all parameters with or without covariates, with the exception of the height parameter without covariates, which showed higher reliability with TBV style preprocessing than with standard preprocessing in the signal without training (see Figure S1).

**Table 6a:**
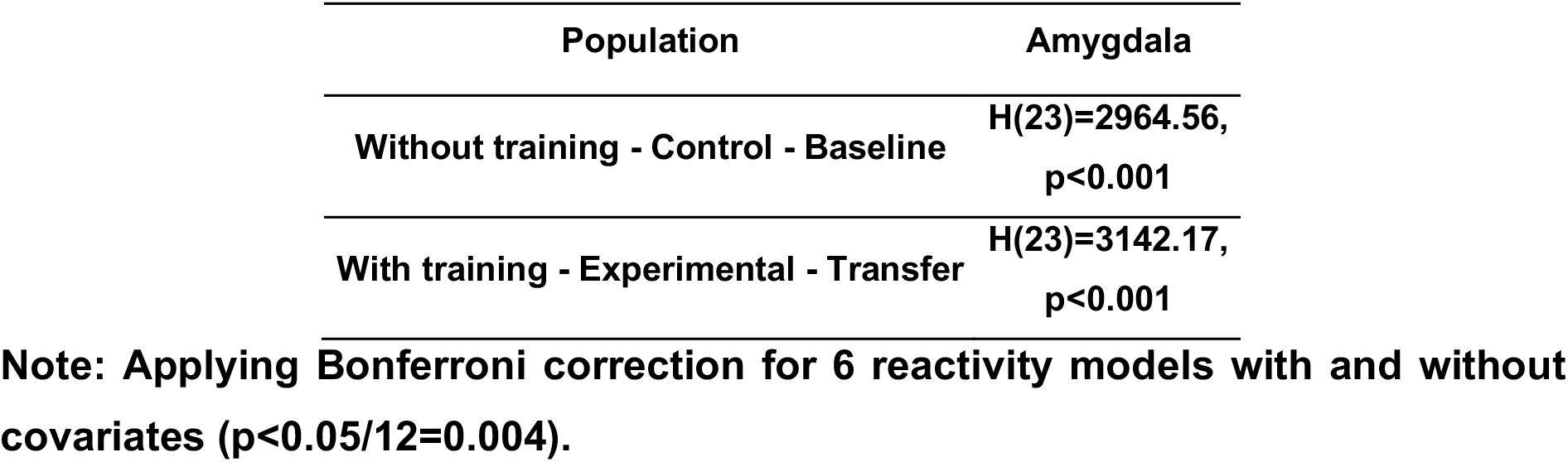
Table of Kruskal Wallis tests’ output for each sample, reactivity model with and without covariates in the left amygdala with Bonferroni correction applied.

| Population | Amygdala |
| --- | --- |
| Without training - Control - Baseline | $H(23)=2964.56,$<br>$p<0.001$ |
| With training - Experimental - Transfer | $H(23)=3142.17,$<br>$p<0.001$ |

**Note: Applying Bonferroni correction for 6 reactivity models with and without covariates ( $p < 0.05/12 = 0.004$ ).**

**Table 6b:** Table of Kruskal Wallis tests’ output for each sample, reactivity model with and without covariates in the left amygdala with Bonferroni correction applied, without rise decay.

| Population | Amygdala |
| --- | --- |
| Without training - Control - Baseline | $H(19)=1397.84,$<br>$p<0.001$ |
| With training - Experimental - Transfer | $H(19)=1702.57,$<br>$p<0.001$ |

**Note: Applying Bonferroni correction for 6 reactivity models with and without covariates ( $p < 0.05/10 = 0.005$ ).**

**Table 7:** Table of number of voxels reaching different reliability thresholds for each sample, preprocessing, and first level parameter with cluster correction applied.

| ROI |  | Amygdala (214 voxels) |  |  |  |  |  |  |  |
| --- | --- | --- | --- | --- | --- | --- | --- | --- | --- |
| Preprocessing |  | BV style |  |  |  | Standard |  |  |  |
| Population | First level model | ICC thresholds |  |  |  | ICC thresholds |  |  |  |
|  |  | 0.4 | 0.6 | 0.7 | 0.75 | 0.4 | 0.6 | 0.7 | 0.75 |
| Without training - Control - Baseline | Canonical amplitude | 0 | 0 | 0 | 0 | 0 | 0 | 0 | 0 |
|  | Amplitude | 52 | 16 | 6 | 2 | 35 | 0 | 0 | 0 |
|  | Area under the curve | 0 | 0 | 0 | 0 | 40 | 0 | 0 | 0 |
|  | Onset-delay | 0 | 0 | 0 | 0 | 0 | 0 | 0 | 0 |
|  | Rise-decay | 0 | 0 | 0 | 0 | 0 | 0 | 0 | 0 |
|  | Height | 78 | 26 | 13 | 13 | 53 | 24 | 9 | 5 |
| With training - Experimental - Transfer | Canonical amplitude | 0 | 0 | 0 | 0 | 0 | 0 | 0 | 0 |
|  | Amplitude | 66 | 4 | 2 | 2 | 42 | 11 | 3 | 2 |
|  | Area under the curve | 0 | 0 | 0 | 0 | 0 | 0 | 0 | 0 |
|  | Onset-delay | 0 | 4 | 4 | 4 | 0 | 5 | 5 | 5 |
|  | Rise-decay | 0 | 0 | 0 | 0 | 0 | 0 | 0 | 0 |
|  | Height | 159 | 81 | 25 | 16 | 73 | 47 | 24 | 21 |

#### 4.2.2. Voxelwise reliability

Some voxelwise ICC values obtained were higher than those computed on the real-time signal covering the entire left amygdala or mean or median ICC values computed over the entire left amygdala (Table 5 vs statistics reported in 4.2.1.1 and Table 6), with some clusters achieving an excellent level of reliability (ICC>.7, see Table 5) for standard and TBV-like preprocessing both for the trained and untrained signals, which did not occur for the region as a whole.

**Table 8:**
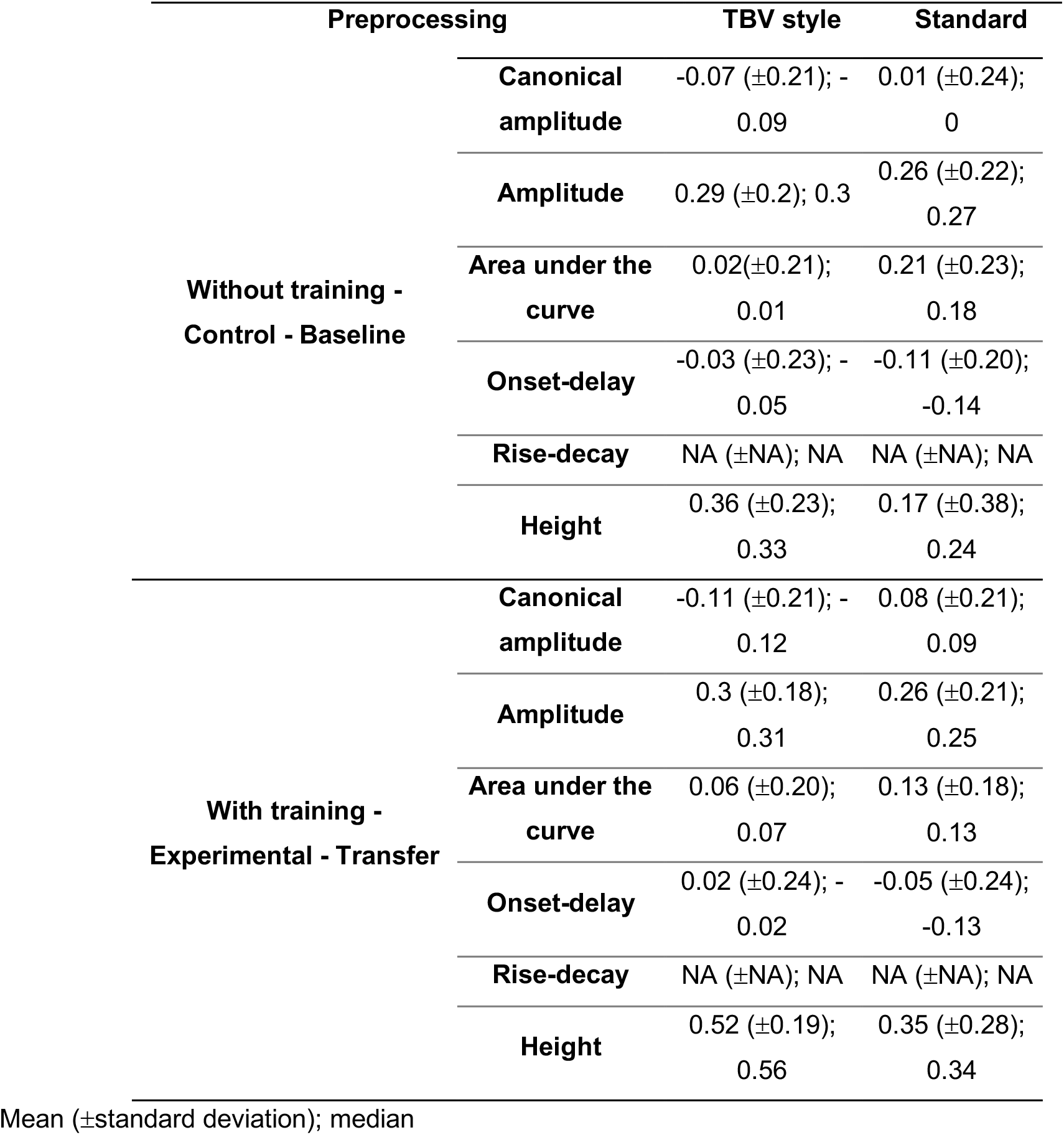
Table of mean, standard deviation and median values of ICCs for each sample, preprocessing, and first level parameter with cluster correction applied.

| Preprocessing | TBV style | Standard |
| --- | --- | --- |

|  |  |  |  |
| --- | --- | --- | --- |
| **Without training -**<br>**Control - Baseline** | **Canonical**<br>**amplitude** | -0.07 ($\pm 0.21$); -<br>0.09 | 0.01 ($\pm 0.24$);<br>0 |
| | **Amplitude** | 0.29 ($\pm 0.2$); 0.3 | 0.26 ($\pm 0.22$);<br>0.27 |
| | **Area under the**<br>**curve** | 0.02($\pm 0.21$);<br>0.01 | 0.21 ($\pm 0.23$);<br>0.18 |
| | **Onset-delay** | -0.03 ($\pm 0.23$); -<br>0.05 | -0.11 ($\pm 0.20$);<br>-0.14 |
| | **Rise-decay** | NA (±NA); NA | NA (±NA); NA |
| | **Height** | 0.36 ($\pm 0.23$);<br>0.33 | 0.17 ($\pm 0.38$);<br>0.24 |
| **With training -**<br>**Experimental - Transfer** | **Canonical**<br>**amplitude** | -0.11 ($\pm 0.21$); -<br>0.12 | 0.08 ($\pm 0.21$);<br>0.09 |
| | **Amplitude** | 0.3 ($\pm 0.18$);<br>0.31 | 0.26 ($\pm 0.21$);<br>0.25 |
| | **Area under the**<br>**curve** | 0.06 ($\pm 0.20$);<br>0.07 | 0.13 ($\pm 0.18$);<br>0.13 |
| | **Onset-delay** | 0.02 ($\pm 0.24$); -<br>0.02 | -0.05 ($\pm 0.24$);<br>-0.13 |
| | **Rise-decay** | NA (±NA); NA | NA (±NA); NA |
| | **Height** | 0.52 ($\pm 0.19$);<br>0.56 | 0.35 ($\pm 0.28$);<br>0.34 |

Mean ( ± standard deviation); median

#### 4.2.3. Clinical and Design Related Measures

##### 4.2.3.1. Amygdala signal

Adding covariates when computing semi-partial correlations over the mean amygdala signal improved descriptively reliability estimates for the signal without training (from sr=0.11 to sr=0.14) as well as parameters tested to fit the signal with training (mean: from sr=0.06 to sr=0.12, onset-delay: from sr=0.14 to sr=0.21, rise-decay: from sr=0.03 to sr=0.14, height: from sr=0.16 to sr=0.29) although in no case we did achieve a fair level of reliability (sr<0.4).

##### 4.2.3.2. Voxelwise signal

The addition of covariates in never resulted in higher average ranks of semipartial correlation distributions on the untrained or trained signal preprocessed with the TBV-like or standard pipeline (Figure S1).

## 5. GENERAL DISCUSSION

As stated in a recent meta-analysis (Elliott et al., 2020), task fMRI reliability is not systematically evaluated and when it is, task-related fMRI measures show poor reliability. Our literature review shows that both prognostic and interventional fMRI studies in MDD, which might otherwise be poised for clinical translation, also do not attend to reliability. We demonstrate that by attending to some fairly simple principles, we can achieve fair to good reliability in a clinical prediction outcome dataset and excellent reliability in a neurofeedback fMRI study dataset (Figure 1). These principles include careful modeling of the BOLD signal, identification of reliable voxels within regions of interest, and calculation of reliability in the population for which translational applications are being considered. Across both datasets, the height parameter from a gamma variate function was the most reliable way to model the BOLD signal, especially among patients with MDD, in some regions of interest, and was, in some combinations of region and population or training condition, more reliable than canonical amplitude, though in other cases the reverse was true (Table 3 and 5 and Figure S1). Consequently, we recommend that researchers explore multiple ways of modeling the BOLD signal, particularly including gamma variate modeling in MDD, before concluding their experiment has low reliability. It may also be helpful for software for real-time analysis of fMRI data to implement alternative, potentially more reliable ways of characterizing BOLD responses in real-time.

Increasingly, functional differentiation of sub-regions of subcortical structures such as the amygdala has been acknowledged as important for fMRI (Balderston, Schultz, Hopkins, & Helmstetter, 2014; Ball et al., 2007; Michely, Rigoli, Rutledge, Hauser, & Dolan, 2020; Roy et al., 2009). The comparison of test-retest reliability estimates obtained on the feedback signal averaged over the whole amygdala versus these same estimates computed voxelwise in the neurofeedback dataset suggest non-uniformity across the amygdala in signal reliability as well; the extent to which these differences explain previous results localizing function to subregions is unclear. Thus, we suggest it may be useful to use a voxel-wise or subregion approach to estimating test-retest reliability. Indeed, this method reveals significantly large clusters of voxels with excellent test-retest reliability in the left amygdala which could be used as masks for neurofeedback targets; our method is easily feasible for new studies. Such excellent reliability, which is a prerequisite for clinical translation, was not attained in our dataset, using the more common computation of median ICCs for each ROI (e.g., as recommended by Caceres et al., 2009) (see Tables 4 and 6).

Contrary to our hypotheses, we did not find that adding covariates to the model, including the scanner on which participants were run and severity, which did change as a function of intervention, improved test-retest reliability in these datasets (Figure S1) in ROI-based or whole-brain analyses (see Figure S2). That said, covariates may still be useful to include in other datasets - we recommend exploring this option further before dismissing their utility.

Reliability did vary by whether the entire sample or only patient’s data were included and by whether or not participants were trained on the task, supporting the potential utility of quantifying reliability on tasks and populations that are relevant for the clinical application intended (Tables 3 and 5 and Figure S1).

There are several limitations of this review and analyses. As we have focused only on MDD, it is unclear whether our conclusions apply transdiagnositically. Improving reliability may require different strategies in other diseases, such as Parkinsons, due to age-related atrophy, increased movement, and differences in neurovascular coupling (Lecrux et al., 2019; Paek et al., 2019). There are many fMRI-based metrics we could have examined, including functional connectivity, volumetric measures, and resting state designs, which all provoke unique considerations for optimizing test-retest reliability, some of which have been explored elsewhere (e.g., Noble et al., 2019). Here, we focused on regional BOLD activity as it is a common feature of prediction and neurofeedback studies. Our published data sets had relatively small number of subjects. This is typical for most clinical fMRI studies, but does raise the concern that the sample is too small and underpowered. Therefore, we strongly encourage the replication of these results and that is also why we have applied these suggestions to two different data sets.

## 6. CONCLUSIONS

To summarize, demonstrating that mechanistic indices are reliable is important before their clinical adoption in prediction or treatment-development. The literature in these areas has implicitly accepted this assumption without testing it. Other non-clinical fMRI studies have shown many of the regions targeted in clinical fMRI studies have fairly low test-retest reliability, which was largely replicated using the most common analytic techniques in our datasets. Yet, we have suggested a few principles that appear to improve the test-retest reliability of the obtained mechanistic signals, have shown their feasibility in two previously published fMRI data sets, and have made code publicly available so that researchers with minimal mathematical and programming knowledge can implement them. Wider adoption of these methods could help to realize the potential of clinical fMRI and could extend to improving psychometrics for other time-varying mechanistic indices.

## Acknowledgements

This work was supported by the National Institutes of Health/ National Institute of Mental Health [grant numbers MH115927, MH106591, MH074807, MH58356, MH69618] and through the Pittsburgh Foundation Emmerling Fund [grant number M2007-0114]. The content is solely the responsibility of the authors and does not necessarily represent the official views of the funding agencies.

