## Supplements for "Importance of test-retest reliability for promoting fMRI based screening and interventions in major depressive disorder"

**Box 1: A Priori Region Definitions**

For a priori regionwise analyses (described in Table S3 below and Figure 2 in the main manuscript) we used the following ROIs:

Three a priori regions for this study (amygdala, DLPFC, rACC) were defined anatomically using anatomical atlases rendered on the Colin-27 Montreal Neurological Institute canonical brain. The fourth (sgACC) was defined meta-analytically, as described below.

**Amygdala**

The full amygdala region, as defined in the AAL atlas was used, as shown below.

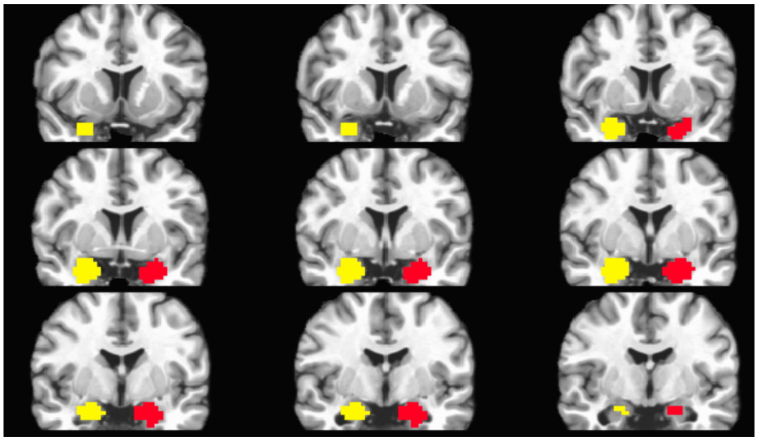

**DLPFC**

The DLPFC was defined as the middle frontal gyrus from AFNI's Talairach (TTatlas) map within 5<Z<37 to yield approximately the lateral BA9/46 region. Depressed and control participants differ on digit sorting as well as the personal relevance task used in this study in this region (Siegle et al., 2007).

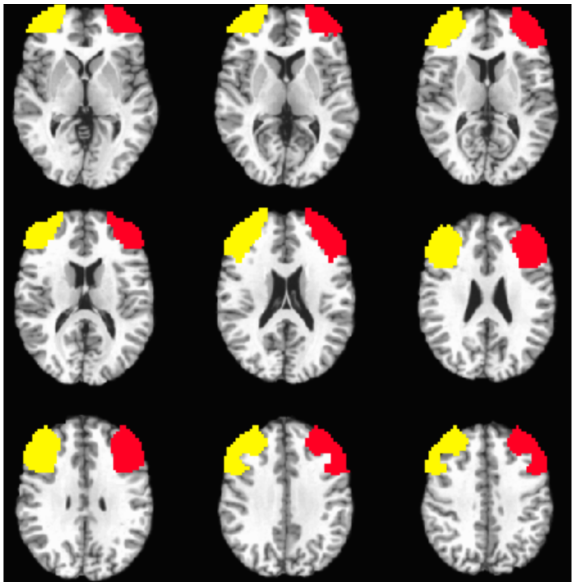

**rACC**

We defined an area of the rostral and subgenual cingulate including BA24 using Afni's Talairach atlas regions for the Anterior Cingulate and Cingulate Gyrus within the range of Y>0 and Z<-15. The rationale for selection of this anatomically defined region is that BA24 in the rostral cingulate consistently predicts response in medication studies using PET (Brannan et al., 2000; Brody et al., 2001; Mayberg et al., 1997), fMRI (Chen et al., 2007; Davidson et al., 2003; Keedwell et al., 2010; Langenecker et al., 2007; Nitschke et al., 2009; Roy et al., 2010), and EEG (Pizzagalli et al., 2001). That said, as the predictive areas have been derived empirically in all of these studies, and do not always overlap, we chose a broader anatomically defined region which does encompass the majority of regions observed in previous studies.

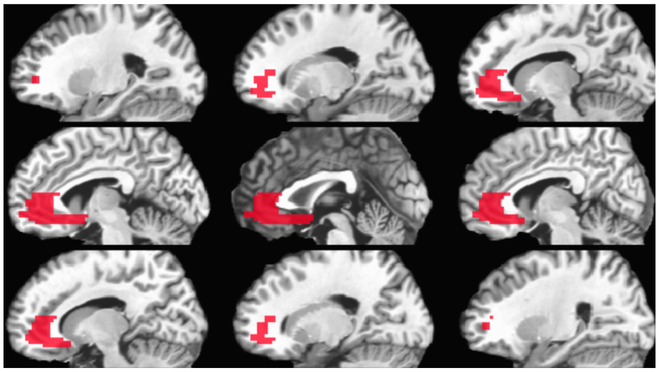

**sgACC**

Our previous studies of Cognitive Behavior Therapy (CBT) have specifically implicated a region of the subgenual cingulate in response to CBT (Siegle et al., 2012). We have recently submitted a meta-analysis of regions which predict response to both CBT and medication in which a region that almost entirely overlaps from that region can also be used to predict response to medication (Strege et al, submitted). We thus used that region. As is always the case, there is some level of arbitrariness to thresholding of meta-analytically defined maps. Our first instinct was to use a conservative thresholding (18 voxels) shown below. That said, we also considered a slightly more liberally thresholded version (33 voxels) as a way of examining the extent to which psychometric characteristics such as reliability could be used to help in defining anatomical masks.

Conservatively thresholded (18 voxels)

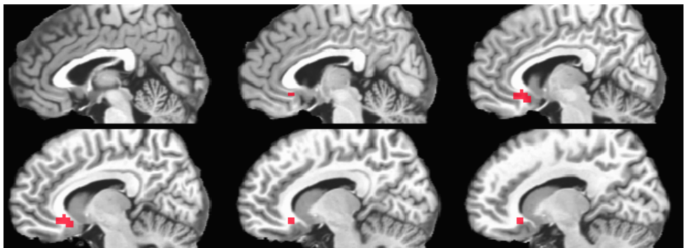

Liberally thresholded (33 voxels)

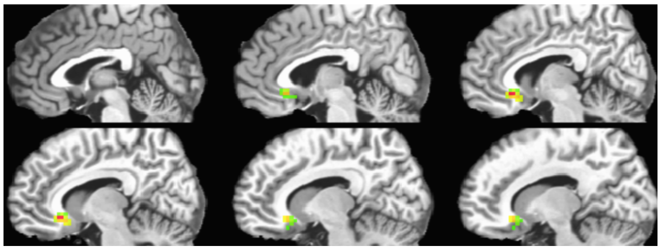

**Box 2: Methodological choice to fit gamma variates**

At the time of the analysis of the feedback signal, two methodological options were possible:

- Either averaging the signal over the blocks of interest and then computing the gamma variates parameters on the mean signal course (choice selected in the main manuscript) [1], or;
- Computing the gamma variates parameters over the signal course in each block and then average the parameters obtained for a run [2].

To make this decision, we tested both option on the left amygdala real-time feedback signal that we extracted from the output of previously used script for real-time preprocessing (Young et al., 2017), see table below.

| Sample | First level gamma variates parameter | Signal averaged over the blocks [1] | Parameter averaged over the run [2] |
|---|---|---|---|
| Signal without training – Baseline in control group | Onset delay | ICC = 0.54 | ICC = 0.03 |
| | Rise decay | ICC = -0.03 | ICC = -0.07 |
| | Height | ICC = 0.47 | ICC = -0.03 |
| Signal with training – Transfer in experimental group | Onset delay | ICC = -0.12 | ICC = -0.13 |
| | Rise decay | ICC = -0.06 | ICC = -0.01 |
| | Height | ICC = -0.12 | ICC = 0.03 |

In conclusion, the second option resulted in poorer test retest reliability results, without this being explained by a poor signal fit. We therefore preferred the first methodology in the main version of the manuscript.

**Box 3: Computation of voxelwise ICCs using different tools**

AFNI's 3dLME and 3dICC functions (Chen et al., 2013) and the FMRELI Matlab toolbox, which is specifically designed for fMRI reliability analyses (Fröhner et al., 2019), support this metric. Both AFNI's 3dLME/3dICC functions and the FMRELI Matlab's toolbox allow computation of type 2 and 3 ICCs. Of these, we have only examined AFNI's 3dLME; it has moderate convergence, which differed by brain region, with well-validated computations implemented in via Matlab (see below). There are also different ways that co-variates can be handled within reliability models. Thus, we note that simply stating that ICC values were computed is unlikely to generalize across analyses – it is at least essential to cite the used software. For the current manuscript, given observed discrepancies, we stayed with a Matlab ICC implementation which we could verify, line for line, with textbook computations, and retreated from the ICC computation to more uniformly accepted methods for computations that necessitated the use of covariates.

Since AFNI's 3dLME function also allows to compute voxel wise ICCs by using a Bayesian approach (-ICCb) to be preferred to an older approach (-ICC), we used that function on the first level canonical amplitude as an example to be able to compare the results obtained from different tools (AFNI and our Matlab function). Using 3dLME, we just added subjects as a random factor, following the example in the online documentation to get an ICC close to type 3, the one that we computed with Matlab. Our Matlab function (icc.m, available from https://github.com/PICANlab/Reliability_toolbox), compute ICC(3,1) as described in Shrout and Fleiss (1979). Then, we computed correlations between ICCs values originated by AFNI and Matlab in all voxels or in ROI:

| Sample | Correlations between ICCs values from Matlab | And from AFNI using Bayesian approach (-ICCb) | And from AFNI using old approach (-ICC) |
|---|---|---|---|
| Patients & controls | On all voxels | r=0.7 | r=0.62 |
| | Voxels within the amygdala | r=0.76 | r=0.71 |
| | Voxels within DLPFC | r=0.87 | r=0.8 |
| | Voxels with rACC | r=0.69 | r=0.49 |
| | Voxels within the sgACC (conservatively thresholded) | r=0.90 | r=0.49 |
| | Voxels within all ROIs | r=0.84 | r=0.77 |
| Patients | On all voxels | r=0.58 | r=0.35 |
| | Voxels within the amygdala | r=0.62 | r=NA |
| | Voxels within DLPFC | r=0.76 | r=0.49 |
| | Voxels with rACC | r=0.31 | r=0.13 |
| | Voxels within the sgACC (conservatively thresholded) | r=0.66 | r=0.27 |
| | Voxels within all ROIs | r=0.63 | r=0.38 |

We wanted to share those results to show that using different tools to get ICC values is likely to give variable results, which also vary based on region.

**Box 4: The particular case of functional localizers**

In many rtfMRI-nf designs, a functional localizer is used to identify the brain region(s) to train (e.g., Linden et al., 2012). Use of functional localizers allows for dynamic adjustment of target areas/networks according to individual patterns, and for evaluation of areas that are most activated by the task at the individual level, but does not guarantee the stability of the signal, nor does it allow a coherent classification among individuals of a group on the basis of the magnitude of activity. Examining the stability of regions activated during a functional localizer task takes into account the variability of interindividual functional specialization, and allows researchers to observe how stable the region/network trained in participants is from one training session to another. Theoretically, regions trained in one session should result in these same regions being more activated during the next session. If the overlap between the target areas from one session to another is weak, it is possible that the training has not been successful. Thus, overlap of functionally defined areas from one session to another is one indication of the reliability of the training procedure. While to our knowledge, no rtfMRI-nf study has reported on the stability of the localizer, this provides neurofeedback researchers an opportunity to examine signal reliability, and we encourage researchers using functional localizers to report the stability/overlap of active voxels selected on different training days at the individual level. In contrast, the field of EEG-neurofeedback, which has broken through to clinical audiences, may have been successful, in part, because it is built on reliability studies of the EEG parameters they subject to neurofeedback (Harmonya et al., 1993; McEvoy et al., 2000; Pollock et al., 1991; Salinsky et al., 1991).

**Box 5: Details regarding datasets used**

Neuroimaging treatment outcome dataset

The sample consisted of 57 patients with major depressive disorder and 34 healthy control participants. Participants were assessed twice, approximately 12-16 weeks apart, during which patients received SSRI medication or CT. Two different scanners were used (both 3T Siemens Trio). The fMRI task was a personal relevance rating task in which participants were asked to indicate the extent to which positive, neutral or negative words were relevant to them or their lives. In this slow event related task, only responses to negative words were analyzed. The University of Pittsburgh Institutional Review Board (IRB) approved this study and all participants gave written consent to participate. Please refer to Siegle et al. (2012a) for a complete description of the design, population, fMRI task, and processing.

Neurofeedback dataset

36 patients (18-55 years), unmedicated, meeting the DSM-IV-TR criteria for MDD and experiencing a major depressive episode participated in a neurofeedback clinical trial during which participants were asked to retrieve positive memories while attempting to increase their hemodynamic activity in the assigned region (illustrated by a thermometer). In this double-blind, placebo-controlled, randomized clinical trial, participants were randomly assigned to receive two sessions of neurofeedback training approximately one week apart, either from the left amygdala or from a parietal control region. The neurofeedback protocol consisted of six runs and no feedback was provided during the first and last runs (baseline and transfer runs, respectively). During each run, participants alternately performed 40-second blocks of rest, happy memories (upregulate condition), and count (backward from 300 by a given number). Feedback was provided only during happy conditions. Imaging was performed using a GE-Discovery MR750 3-T scanner equipped with a custom rtfMRI neurofeedback system. The Western Institutional Review Board approved this research protocol that was registered on ClinicalTrials.gov and participants gave consent to participate in the study. Please refer to Young, Siegle, et al. (2017) for a complete description of the design, population and fMRI task.

**Figure S1: Average group ranks and confidence intervals for semi partial correlations distributions for each model in each ROI in each data set.**

| Siegle et al. (2012) data set | | |
|---|---|---|
| **ROI** | **Controls & patients** | **Patients** |
| Amygdala | 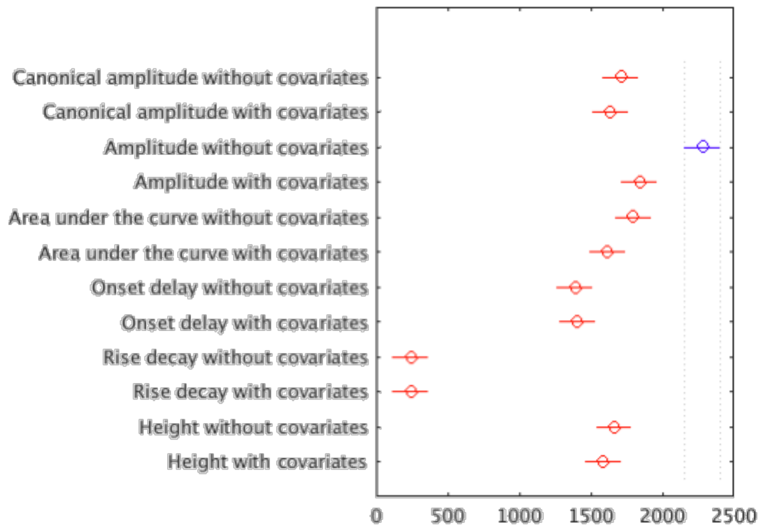 | 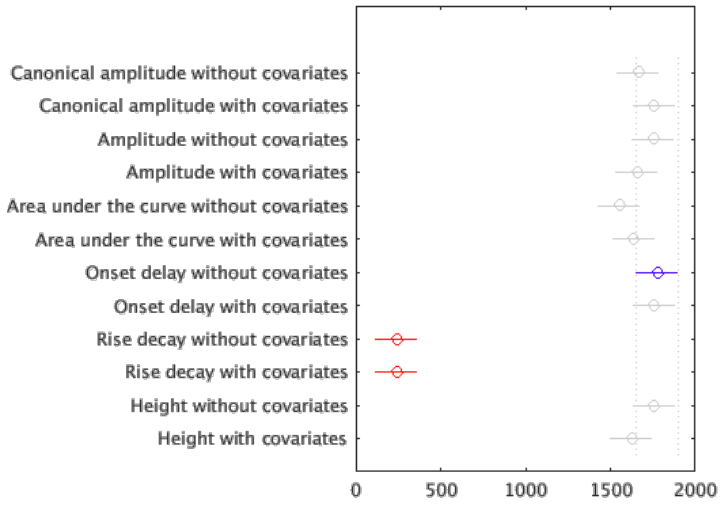 |

DLPFC

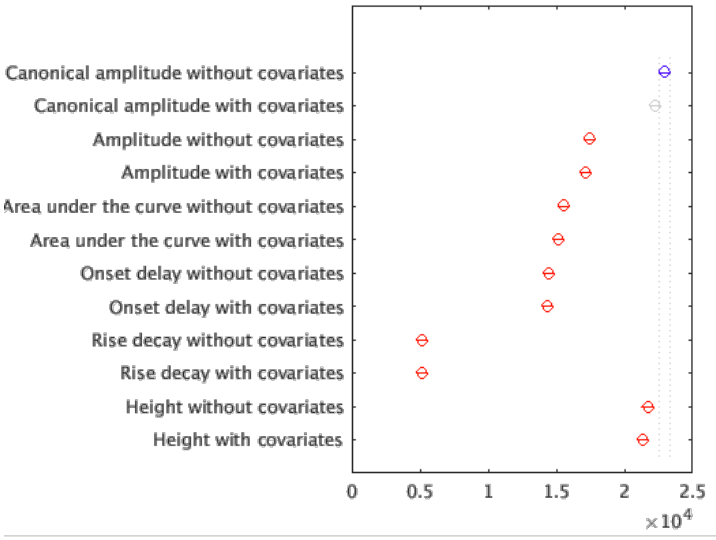
Canonical amplitude without covariates
Canonical amplitude with covariates
Amplitude without covariates
Amplitude with covariates
Area under the curve without covariates
Area under the curve with covariates
Onset delay without covariates
Onset delay with covariates
Rise decay without covariates
Rise decay with covariates
Height without covariates
Height with covariates
0
0.5
1
1.5
2
2.5
$\times 10^4$

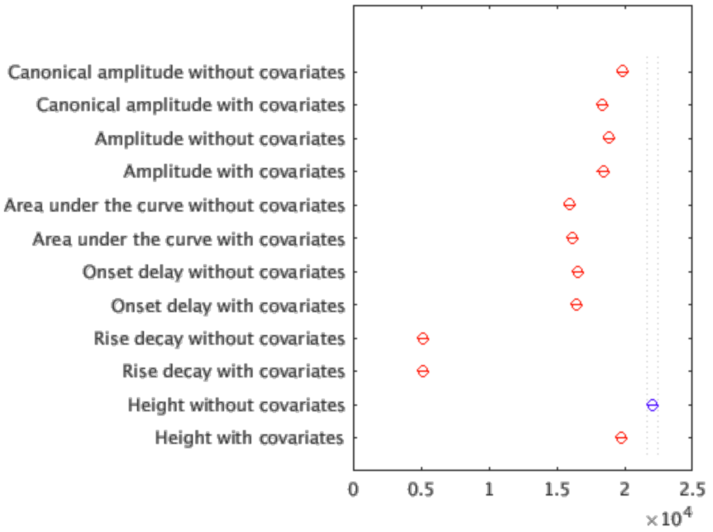
Canonical amplitude without covariates
Canonical amplitude with covariates
Amplitude without covariates
Amplitude with covariates
Area under the curve without covariates
Area under the curve with covariates
Onset delay without covariates
Onset delay with covariates
Rise decay without covariates
Rise decay with covariates
Height without covariates
Height with covariates
0
0.5
1
1.5
2
2.5
$\times 10^4$

rACC

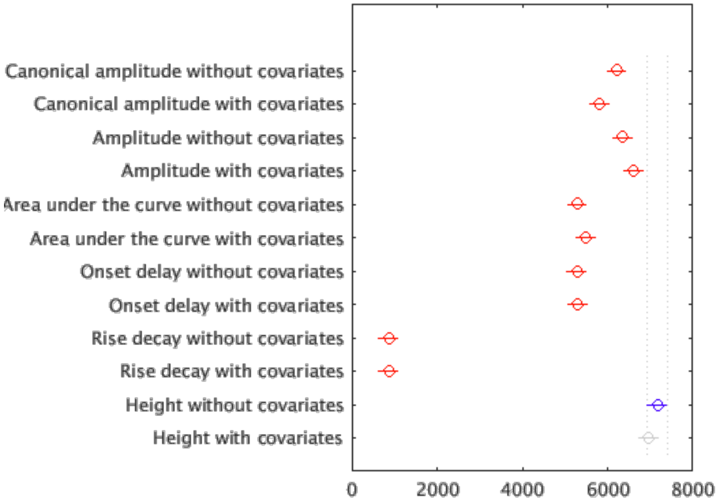
Canonical amplitude without covariates
Canonical amplitude with covariates
Amplitude without covariates
Amplitude with covariates
Area under the curve without covariates
Area under the curve with covariates
Onset delay without covariates
Onset delay with covariates
Rise decay without covariates
Rise decay with covariates
Height without covariates
Height with covariates
0
2000
4000
6000
8000

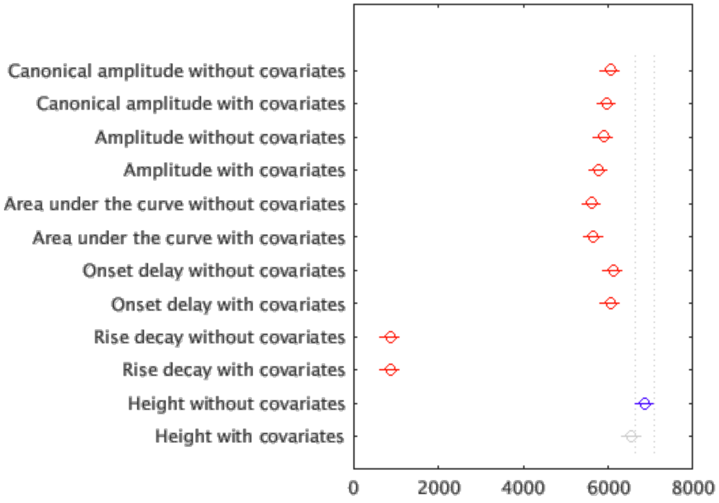
Canonical amplitude without covariates
Canonical amplitude with covariates
Amplitude without covariates
Amplitude with covariates
Area under the curve without covariates
Area under the curve with covariates
Onset delay without covariates
Onset delay with covariates
Rise decay without covariates
Rise decay with covariates
Height without covariates
Height with covariates
0
2000
4000
6000
8000

| | | |
|---|---|---|
| sgACC liberally thresholded | 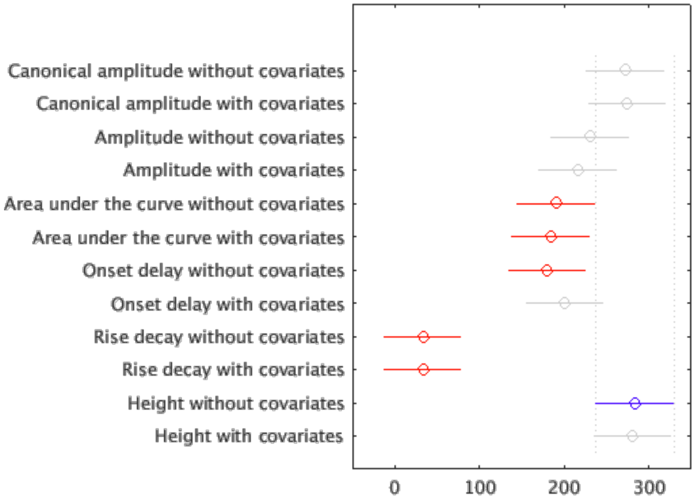 | 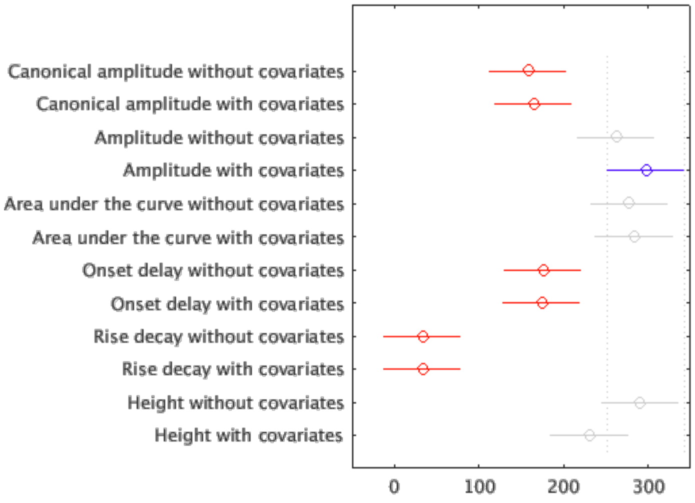 |
| sgACC conservativel y thresholded | 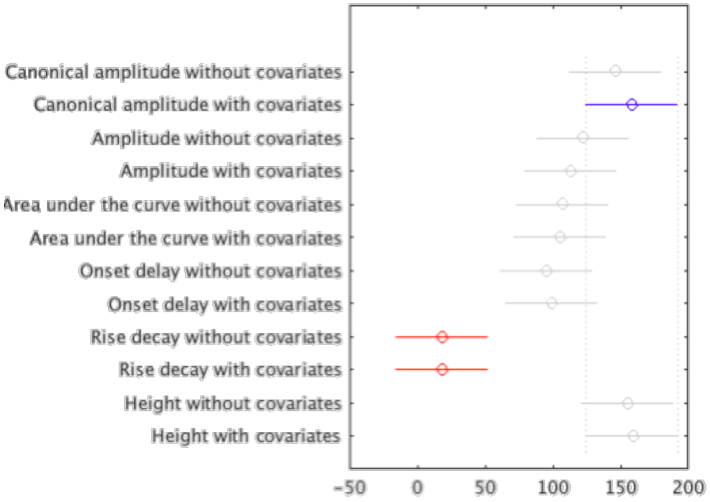 | 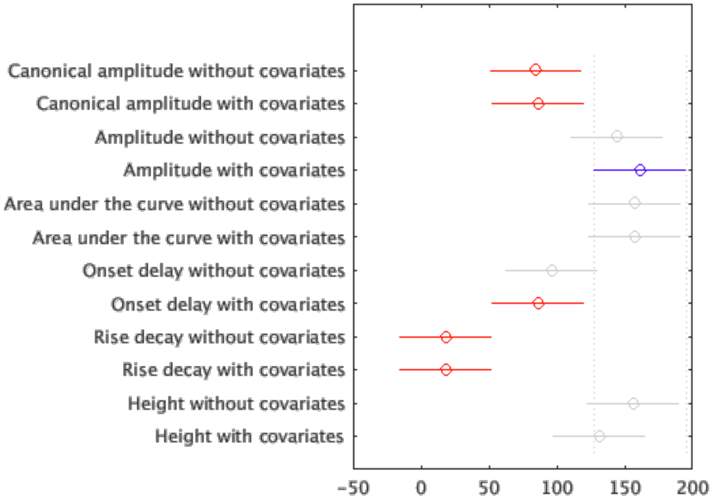 |

### Young et al. (2017b) data set

<table>
<tr><th>ROI</th><th>Signal without training</th><th>Trained signal</th></tr>
<tr><td>Amygdala</td><td>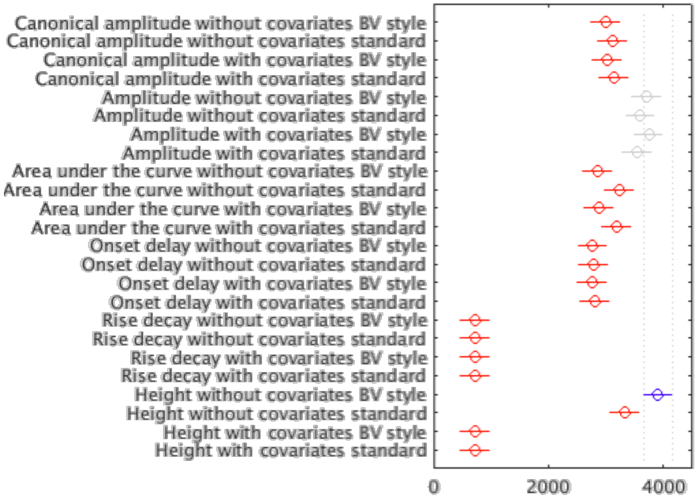
</td><td>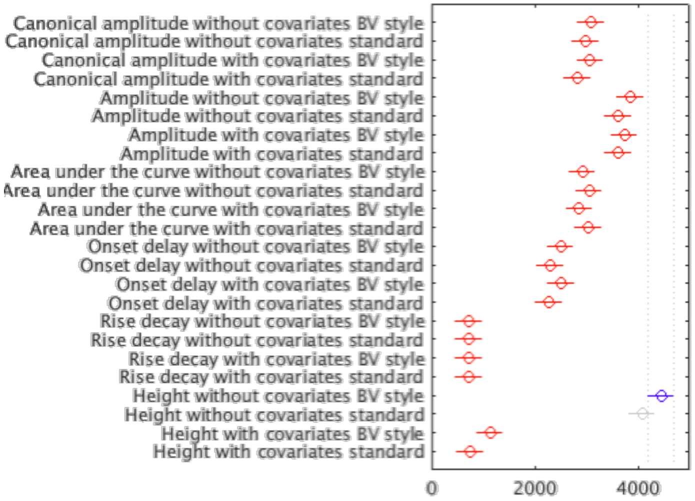
</td></tr>
</table>

**Figure S2: Voxelwise benefit from adding covariates, rtfMRI-nf dataset (fMRI activation task dataset)**

Positive semi partial correlation difference computed with covariates versus without covariate in each voxel with the height gamma variates parameter A. For controls and patients. B. For patients only.

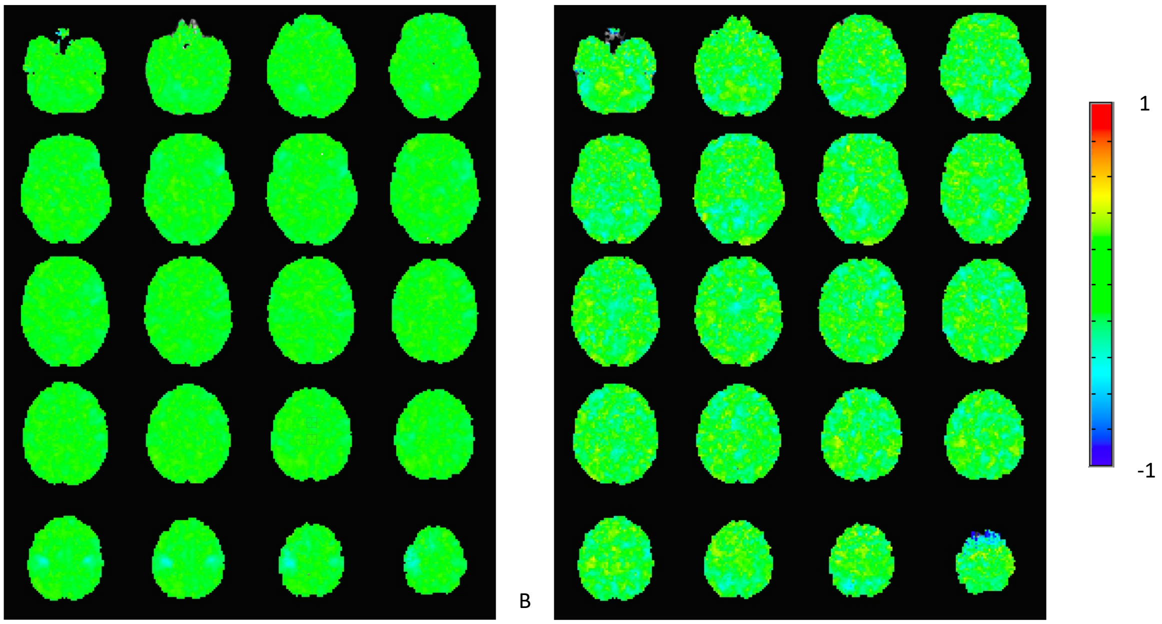

A

**Table S1: Table of rescaled average group ranks of semi partial correlations for each sample, first level parameter, and ROI, with and without covariates (fMRI activation task dataset)**

| Population | First level model | Covariates | Amygdala | DLPFC | rACC | sgACC liberally thresholded | sgACC conservatively thresholded |
|---|---|---|---|---|---|---|---|
| Controls & patients | Canonical amplitude | Without | 7.07 | 8.6 | 7.2 | 8.16 | 8.25 |
| | | With | 6.77 | 8.31 | 6.73 | 8.82 | 8.32 |
| | Amplitude | Without | 9.44 | 6.51 | 7.36 | 6.81 | 6.99 |
| | | With | 7.62 | 6.42 | 7.65 | 6.33 | 6.56 |
| | Area under the curve | Without | 7.44 | 5.81 | 6.12 | 5.95 | 5.79 |
| | | With | 6.68 | 5.68 | 6.36 | 5.87 | 5.6 |
| | Onset delay | Without | 5.75 | 5.39 | 6.1 | 5.28 | 5.45 |
| | | With | 5.82 | 5.37 | 6.13 | 5.53 | 6.09 |
| | Rise decay | Without | 1 | 1.91 | 1 | 1.03 | 1.02 |
| | | With | 1 | 1.91 | 1 | 1.03 | 1.02 |
| | Height | Without | 6.88 | 8.11 | 8.29 | 8.64 | 8.58 |
| | | With | 6.56 | 7.97 | 8.06 | 8.88 | 8.51 |
| Patients | Canonical amplitude | Without | 6.89 | 7.41 | 7 | 4.72 | 4.79 |
| | | With | 7.27 | 6.87 | 6.89 | 4.83 | 4.99 |
| | Amplitude | Without | 7.24 | 7.05 | 6.81 | 8.02 | 7.94 |
| | | With | 6.85 | 6.88 | 6.68 | 8.98 | 9.01 |
| | Area under the curve | Without | 6.42 | 5.97 | 6.49 | 8.74 | 8.41 |
| | | With | 6.77 | 6.05 | 6.54 | 8.74 | 8.59 |
| | Onset delay | Without | 7.34 | 6.18 | 7.07 | 5.38 | 5.32 |
| | | With | 7.26 | 6.14 | 7 | 4.83 | 5.29 |
| | Rise decay | Without | 1 | 1.91 | 1 | 1.03 | 1.02 |
| | | With | 1 | 1.91 | 1 | 1.03 | 1.02 |
| | Height | Without | 7.26 | 8.24 | 7.94 | 8.71 | 8.8 |
| | | With | 6.72 | 7.39 | 7.58 | 7.33 | 7 |

**Table S2: Table of rescaled average group ranks of semi partial correlations for each sample, preprocessing, first level parameter, and ROI, with and without covariates (rtfMRI-nf dataset)**

| Preprocessing | | | BV style | Standard |
|---|---|---|---|---|
| Without training Control - Baseline | Canonical amplitude | Without | 7.01 | 7.29 |
| | | With | 7.05 | 7.34 |
| | Amplitude | Without | 8.66 | 8.41 |
| | | With | 8.75 | 8.26 |
| | Area under the curve | Without | 6.65 | 7.56 |
| | | With | 6.71 | 7.42 |
| | Onset delay | Without | 6.47 | 6.49 |
| | | With | 6.46 | 6.56 |
| | Rise decay | Without | 1.67 | 1.67 |
| | | With | 1.67 | 1.67 |
| | Height | Without | 9.13 | 7.77 |
| | | With | 1.67 | 1.67 |
| With training Experimental Transfer | Canonical amplitude | Without | 7.24 | 6.99 |
| | | With | 7.19 | 6.62 |
| | Amplitude | Without | 9 | 8.46 |
| | | With | 8.73 | 6.46 |
| | Area under the curve | Without | 6.83 | 7.14 |
| | | With | 6.7 | 7.1 |
| | Onset delay | Without | 5.85 | 5.38 |
| | | With | 5.89 | 5.32 |
| | Rise decay | Without | 1.71 | 1.71 |
| | | With | 1.71 | 1.71 |
| | Height | Without | 10.39 | 9.53 |
| | | With | 2.65 | 1.75 |

**Table S3: Table of number of contiguous voxels for used cluster correction and p values associated for each reliability threshold, ROI, group for treatment outcome data set**

| Population | First level model | Reliability level | Amygdala | DLPFC | rACC | sgACC liberally thresholded | sgACC conservatively thresholded |
|---|---|---|---|---|---|---|---|
| Controls & patients (N=91) | Canonical amplitude | **0.4** | 1 | 1.8 | 1.1 | - | - |
| | | **0.6** | 1 | 1 | 1 | - | - |
| | | **0.7** | 1 | 1 | 1 | - | - |
| | | **0.75** | 1 | 1 | 1 | - | - |
| | Amplitude | **0.4** | 2 | 1.6 | 1.1 | - | - |
| | | **0.6** | 1 | 1 | 1 | - | - |
| | | **0.7** | 1 | 1 | 1 | - | - |
| | | **0.75** | 1 | 1 | 1 | - | - |
| | Area under the curve | **0.4** | 1 | 1.7 | 1.1 | - | - |
| | | **0.6** | 1 | 1 | 1 | - | - |
| | | **0.7** | 1 | 1 | 1 | - | - |
| | | **0.75** | 1 | 1 | 1 | - | - |
| | Onset delay | **0.4** | 1 | 1.3 | 1.1 | - | - |
| | | **0.6** | 1 | 1 | 1 | - | - |
| | | **0.7** | 1 | 1 | 1 | - | - |
| | | **0.75** | 1 | 1 | 1 | - | - |
| | Rise decay | **0.4** | - | - | - | - | - |
| | | **0.6** | - | - | - | - | - |
| | | **0.7** | - | - | - | - | - |
| | | **0.75** | - | - | - | - | - |
| | Height | **0.4** | 1 | 1.8 | 1.1 | - | - |
| | | **0.6** | 1 | 1 | 1 | - | - |
| | | **0.7** | 1 | 1 | 1 | - | - |
| | | **0.75** | 1 | 1 | 1 | - | - |
| Patients | Canonical amplitude | **0.4** | 2.6 | 6.7 | 4 | - | - |
| | | **0.6** | 1 | 1 | 1 | - | - |
| | | **0.7** | 1 | 1 | 1 | - | - |
| | | **0.75** | 1 | 1 | 1 | - | - |
| | Amplitude | **0.4** | 2 | 7.2 | 3.7 | - | - |
| | | **0.6** | 1 | 1 | 1 | - | - |

| | | | | | | |
|---|---|---|---|---|---|---|
| | **0.7** | 1 | 1 | 1 | - | - |
| | **0.75** | 1 | 1 | 1 | - | - |
| **Area under the curve** | **0.4** | 2.2 | 9.7 | 3.9 | - | - |
| | **0.6** | 1 | 1 | 1 | - | - |
| | **0.7** | 1 | 1 | 1 | - | - |
| | **0.75** | 1 | 1 | 1 | - | - |
| **Onset delay** | **0.4** | 1.6 | 2.6 | 2 | - | - |
| | **0.6** | 1 | 1 | 1 | - | - |
| | **0.7** | 1 | 1 | 1 | - | - |
| | **0.75** | 1 | 1 | 1 | - | - |
| **Rise decay** | **0.4** | - | - | - | - | - |
| | **0.6** | - | - | - | - | - |
| | **0.7** | - | - | - | - | - |
| | **0.75** | - | - | - | - | - |
| **Height** | **0.4** | 2.0 | 7.5 | 3.9 | - | - |
| | **0.6** | 1 | 1 | 1 | - | - |
| | **0.7** | 1 | 1 | 1 | - | - |
| | **0.75** | 1 | 1 | 1 | - | - |

Notes: After getting different numbers of voxels to use for cluster correction while running 3dClustsim two times in a row with a standard number of 2,000 simulations, we increased the number of iterations to 10,000 to obtain more stability in our results.

For 91 participants in whole sample group r=.4 yields p=.100025.

For 91 participants in whole sample group r=.6 yields p=.008479.

For 91 participants in whole sample group r=.7 yields p=.001219.

For 91 participants in whole sample group r=.75 yields p=.000338.

For 57 patients in patients group r=.4 yields p=.12475.

For 57 patients in patients group r=.6 yields p=.014007.

For 57 patients in patients group r=.7 yields p=.002535.

For 57 patients in patients group r=.75 yields p=.00082.

AFNI function 3dClustSim was not able to compute the number of voxels necessary for cluster correction with the masks of sgACC liberally and conservatively thresholded because of their small size (respectively, 33 and 18 voxels) so in this case, no correction was applied and on the first parameter rise decay rate in every ROI because data were badly fitted so there was no variance in the ICC values.

AFNI 3dFWHMx function for the residual ICC map with the first level parameter amplitude in the whole sample had a calculation error using ACF method with the original amygdala ROI so we dilated this ROI of one voxel in this only particular case.

**Table S4: Table of number of contiguous voxels for used cluster correction and p values associated for each reliability threshold, ROI, group and preprocessing for neurofeedback dataset**

| Preprocessing | | | BV style | Standard |
|---|---|---|---|---|
| Without training Control - Baseline | Canonical amplitude | **0.4** | 30.3 | 31.3 |
| | | **0.6** | 4.1 | 4.9 |
| | | **0.7** | 2.1 | 2.2 |
| | | **0.75** | 1.4 | 1.5 |
| | Amplitude | **0.4** | 30.6 | 30.5 |
| | | **0.6** | 4.3 | 4.4 |
| | | **0.7** | 2.1 | 2.1 |
| | | **0.75** | 1.4 | 1.4 |
| | Area under the curve | **0.4** | 35.6 | 36 |
| | | **0.6** | 6.6 | 6.2 |
| | | **0.7** | 2.8 | 2.5 |
| | | **0.75** | 1.7 | 1.5 |
| | Onset delay | **0.4** | 27.8 | 28.7 |
| | | **0.6** | 3.1 | 3.3 |
| | | **0.7** | 1.7 | 1.7 |
| | | **0.75** | 1.3 | 1.3 |
| | Rise decay | **0.4** | - | - |
| | | **0.6** | - | - |
| | | **0.7** | - | - |
| | | **0.75** | - | - |
| | Height | **0.4** | 28.2 | 28.3 |
| | | **0.6** | 3.3 | 3.4 |
| | | **0.7** | 1.7 | 1.8 |
| | | **0.75** | 1.3 | 1.3 |
| With training Experimental Transfer | Canonical amplitude | **0.4** | 28.3 | 29.4 |
| | | **0.6** | 4.1 | 4.7 |
| | | **0.7** | 1.7 | 1.8 |
| | | **0.75** | 1.1 | 1.1 |
| | Amplitude | **0.4** | 26.3 | 26.2 |
| | | **0.6** | 3.6 | 4.3 |

| | | | |
|---|---|---|---|
| | **0.7** | 1.7 | 1.8 |
| | **0.75** | 1.1 | 1.1 |
| **Area under the curve** | **0.4** | 24.7 | 30 |
| | **0.6** | 3.9 | 5.2 |
| | **0.7** | 1.8 | 1.9 |
| | **0.75** | 1.1 | 1.1 |
| **Onset delay** | **0.4** | 20 | 19.6 |
| | **0.6** | 2.4 | 2.4 |
| | **0.7** | 1.4 | 1.4 |
| | **0.75** | 1.1 | 1.1 |
| **Rise decay** | **0.4** | - | - |
| | **0.6** | - | - |
| | **0.7** | - | - |
| | **0.75** | - | - |
| **Height** | **0.4** | 21.1 | 20.5 |
| | **0.6** | 2.8 | 2.7 |
| | **0.7** | 1.5 | 1.4 |
| | **0.75** | 1.1 | 1.1 |

Notes: After getting different numbers of voxels to use for cluster correction while running 3dClustsim two times in a row with a standard number of 2,000 simulations, we increased the number of iterations to 10,000 to obtain more stability in our results.

For 18 patients in experimental group r=.4 yields p=.100025.

For 18 patients in experimental group r=.6 yields p=.008479.

For 18 patients in experimental group r=.7 yields p=.001219.

For 18 patients in experimental group r=.75 yields p=.000338.

For 16 patients in control group r=.4 yields p=.12475.

For 16 patients in control group r=.6 yields p=.014007.

For 16 patients in control group r=.7 yields p=.002535.

For 16 patients in control group r=.75 yields p=.00082.

AFNI function 3dClustSim was not able to compute the number of voxels necessary for cluster correction on the first parameter rise decay rate because data were badly fitted so there was no variance in the ICC values.
